# Hierarchical Markov Random Field model captures spatial dependency in gene expression, demonstrating regulation via the 3D genome

**DOI:** 10.1101/2019.12.16.878371

**Authors:** Naihui Zhou, Iddo Friedberg, Mark S. Kaiser

## Abstract

HiC technology has revealed many details about the eukaryotic genome’s complex 3D architecture. It has been shown that the genome is separated into organizational structures which are associated with gene expression. However, to the best of our knowledge, no studies have quantitatively measured the level of gene expression in the context of the 3D genome.

Here we present a novel model that integrates data from RNA-seq and HiC experiments, and determines how much of the variation in gene expression can be accounted for by the genes’ spatial locations. We used Poisson hierarchical Markov Random Field (PhiMRF), to estimate the level of spatial dependency among protein-coding genes in two different human cell lines. The inference of PhiMRF follows a Bayesian framework, and we introduce the Spatial Interaction Estimate (SIE) to measure the strength of spatial dependency in gene expression.

We find that the quantitative expression of genes in some chromosomes show meaningful positive intra-chromosomal spatial dependency. Interestingly, the spatial dependency is much stronger than the dependency based on linear gene neighborhoods, suggesting that 3D chromosome structures such as chromatin loops and Topologically Associating Domains (TADs) are strongly associated with gene expression levels. In some chromosomes the spatial dependency in gene expression is only detectable when the spatial neighborhoods are confined within TADs, suggesting TAD boundaries serve as insulating barriers for spatial gene regulation in the genome. We also report high inter-chromosomal spatial correlations in the majority of chromosome pairs, as well as the whole genome. Some functional groups of genes show strong spatial dependency in gene expression as well, providing new insights into the regulation mechanisms of these molecular functions. This study both confirms and quantifies widespread spatial correlation in gene expression. We propose that, with the growing influx of HiC data complementing gene expression data, the use of spatial dependence should be an integral part of the toolkit in the computational analysis of the relationship between chromosome structure and gene expression.

## 1 Introduction

The 3D genome organization plays an important role in gene expression through various mechanisms [1, 2, 3, 4]. Of special interest is how genes in close spatial proximity coordinate expression. Several molecular models involving different organizational hierarchies have been proposed to explain this phenomenon [4]. One such hypothesis is that of *transcription factories*, where RNA polymerase II is significantly enriched to allow efficient transcription of multiple genes at the same foci [5, 6, 7].

Another molecular model for spatial gene clusters hypothesizes that the spatial cluster are brought together to allow their promoters to interact with enhancers [8, 9]. The TNF*α*-induced multigene complex regulated by NF-*κ*B is disrupted once the chromatin loops for the complex are cleaved [10], potentially explained by the fact that these genes are dependent on NF-*κ*B-responsive enhancers [11]. Indeed, genes sharing common regulatory elements through a promoter interaction network are spatially co-localized with correlated expression levels [12].

Many of the aforementioned studies are made possible following the advent of Chromosomal Conformation Capture (3C) [13] and subsequent 4C [14, 15], HiC [16, 11, 17] and Capture HiC [12, 9] technologies. These advances allow for a global overview of the genomic architecture instead of individual loci. HiC has enabled or confirmed discoveries of a hierarchy of organizational structures, from Topologically Associating Domains (TAD) [18], to A/B compartments [16] and Chromosomal Territories (CT) [19]. There is a known general association between these structures and gene expression [3, 20]. For example, the A compartments of the genome are more gene dense and are more actively transcribing[16]. Moreover, disrupting TAD boundaries may result in disruption in expression [21].

Several whole-genome computational studies have attempted to untangle the relationship between gene co-expression and their spatial organization. Inter-chromosomal co-expression is significantly enhanced for genes with spatial contacts in yeast [22], and gene interaction networks can predict co-expression well [23]. Gene pair functional similarity is also correlated with spatial distance [24]. However, these studies represented the expression as a pairwise property, either co-expression or co-functionality, but do not model actual expression levels. Even though these studies confirm that there is a general trend of co-localization for co-regulated, co-expressed or co-functional genes, none of them provide a probabilistic model for gene expression levels within a genome. At the same time, many routine analyses in RNA-seq data, rely heavily on a good probabilistic model of gene expression [25, 26, 27]. The between-sample probability that these methods model is dependent on the between-gene variation in the genome, which can be further modeled given more explanatory data, such as spatial location. This added correlation between the genes in one sample could improve the performance of differential expression analyses tools.

We have developed a probabilistic model, **PhiMRF** (**P**oisson **hi**erarchical **M**arkov **R**andom **F**ield) that integrates spatial location and gene expression. We further introduce the *Spatial Interaction Estimate*, or SIE, a measure whose value indicates the strength of spatial dependency of gene expression. SIE can be thought of as analogous to the more familiar regression slope (*β*). While the slope measures the correlation between two variables, SIE measures the strength of spatial dependency of gene expression. Using PhiMRF, we quantify spatial dependency of gene expression for all chromosomes, as well as for select functional gene groups. The ability to quantify spatial dependence is a considerable advance in understanding the spatial component in the regulation of gene expression. PhiMRF can be used to explore the 3D regulation mechanism of any gene group of interest. With the advent of HiC data regularly added to genomic and transcriptomic data, it is expected that interrogating novel mechanisms of regulation based on the 3D genome structure will become possible and widely used. The statistical framework developed int this work, and PhiMRF provide the means to do so.

## 2 Results

### 2.1 Hierarchical Markov Random Field Model

PhiMRF attributes the variation in gene expression (observed in *k* replicates) to the genes’ 3D locations (observed via HiC experiemnts) using an autoregression-based model (Figure 1). Autoregressive (AR) models are used to explain variation for data observed in spatial or temporal settings by modeling the distribution of an observation as dependent on its past (time series) or on its spatial neighbors. We extract a neighborhood network for gene locations in 3D space from HiC data (Figure 1), where a gene is mapped to all of its overlapping loci (Supplementary Figure S1), and its spatial proximity to another gene is calculated as the summary of all of the loci pairs the two gene covers (Methods).

**Figure 1:**
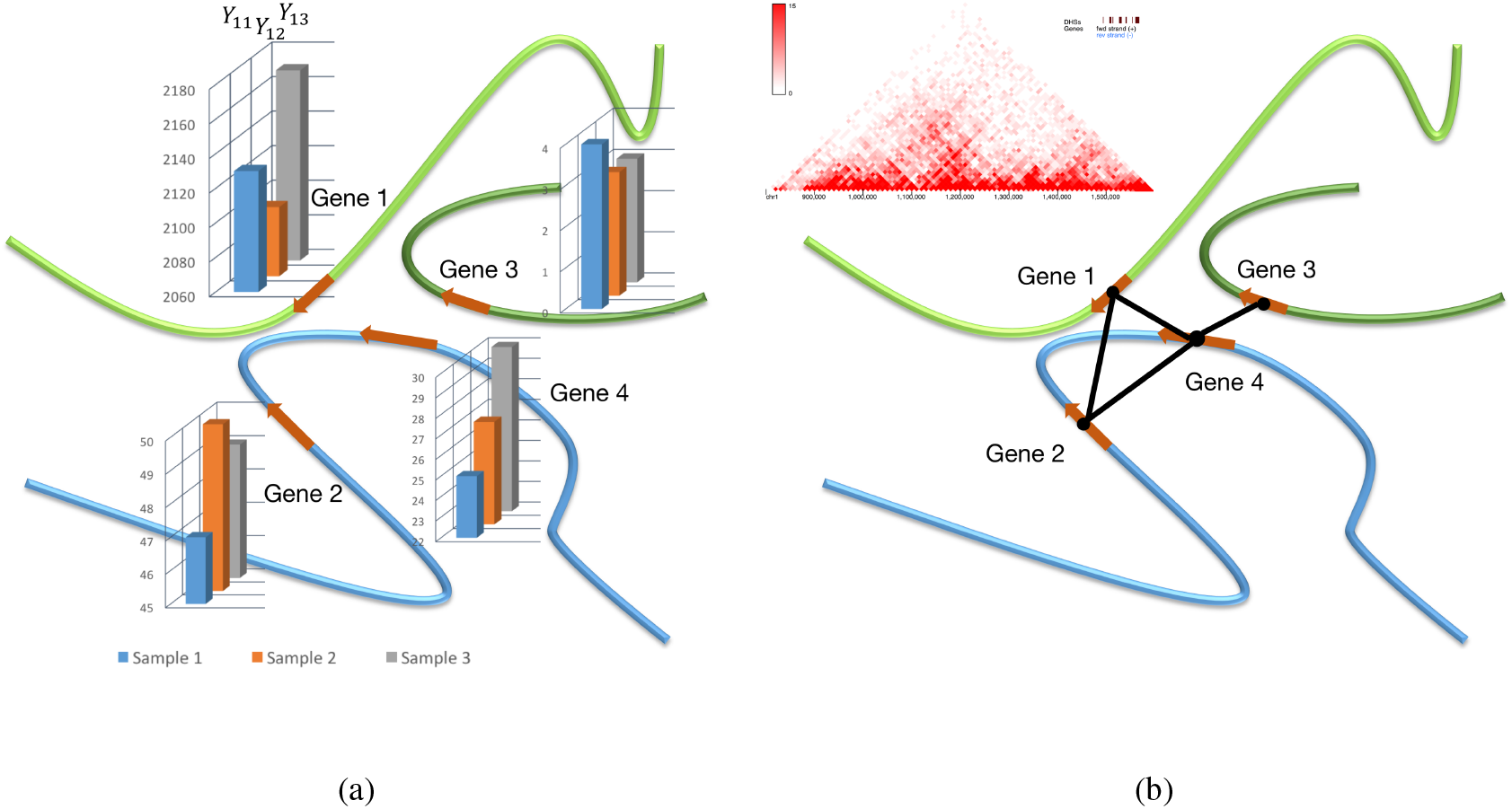
Overall Scheme of the PhiMRF model applied to RNA-seq and HiC data. **(a)** Replicates of RNA-seq quantification can be observed at each gene. **(b)** A spatial gene network is inferred from HiC data. Each gene is treated as a node in the network. An edge exists between two genes if the spatial interaction frequency between loci overlapping with the two genes is higher than a threshold. See Methods for detailed description of the network inference. The triangular HiC heat map is generated using the 3D genome browser [28] with data from Rao *et al* [17].

Our Poisson Hierarchical Markov Random Field (PhiMRF) model is briefly described below (For a detailed description, see Methods and Supplementary Methods). Let *Y*_*ik*_ be the random variable connected with the RNA-seq count for gene *i* (located at location *s*_*i*_) from sample *k*, *i* = 1, 2,…, *n*; *k* = 1, 2,…, *M*. *Y*_*ik*_ is modelled with a Poisson distribution [29], with its parameter *λ*_*i*_, i.e. *Y*_*ik*_ ∼ Poisson(*λ*_*i*_). Let *w*_*i*_ = *log*(*λ*_*i*_). We **conditionally** specify the distribution for *w*_*i*_ as,

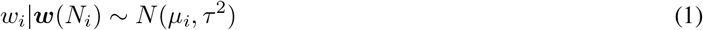

where *N*_*i*_ is the set of locations neighboring *s*_*i*_: *N*_*i*_ = {*s*_*j*_: *s*_*j*_ is a neighbor of *s*_*i*_} and ***w***(*N*_*i*_) = {*w*_*j*_: *s*_*j*_ is a neighbor of *s*_*i*_}. Equation 1 is a conditionally specified model, its mean is further modelled as in Equation 2:

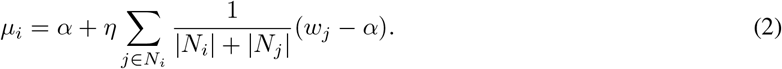

Throughout this study, the posterior distribution of *η* helps us to understand the strength of the spatial dependency. The main properties of the posterior *η* distribution that are the mean 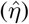 and the 95% credible interval, which is obtained as the 2.5% and 97.5% quantiles of the simulated posterior distribution. We will refer to the estimated posterior mean of *η* 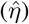 as the **Spatial Interaction Estimate (SIE)**. If the 95% posterior credible interval for *η* does not contain 0, we say that there is meaningful spatial dependency. The two other unknown parameters in this model are *α* and *τ*^2^, where *α* is connected with a basal expression rate for all genes. The parameter *τ*^2^ is connected with the conditional Gaussian variance, which accounts for any remaining variance in gene expression within a sample. The same properties (mean, 2.5% and 97.5% quantiles) are used to summarize their posterior distributions.

In summary, PhiMRF models gene expression with a conditional Poisson-lognormal mixture, and a autoregressive model with a parameter *η* that is connected with spatial dependency. We applied Bayesian inference that allowed us the simulate from the posterior distributions of *η*, resulting in the Spatial Interaction Estimate (SIE) that symbolizes the strength of the spatial dependency.

### 2.2 Intra-chromosomal dependency

#### Within each chromosome

We ran PhiMRF on all genes in each of the 23 human chromosomes (Y chromosome excluded) in the IMR90 cell, with the spatial gene networks inferred from intra-chromosomal HiC data with 10kb resolution [17]. We also implemented a linear baseline for each chromosome. The baseline takes the same gene expression data for each gene but the spatial network is simply inferred from genes within 10k base pairs of each other in the linear chromosome. By comparing our data with this baseline dataset, we can observe whether the long-range, non-linear interactions do play a role in gene expression. We found strong evidence of positive spatial dependency in eleven chromosomes: 1, 4, 5, 6, 8, 9, 12, 19, 20, 21 and X, with SIEs higher than the linear baseline (Figure 2a, Supplementary Tables S1 and S2). This suggested genes in these chromosomes are co-dependent on their neighbors when it comes to gene expression, proving that genes that are spatially close have coordinated expression patterns on a global scale. We noticed that although in all these eleven chromosomes the 95% credible interval of *η* is greater than zero, their ranges vary greatly. Although these SIEs are all larger than their linear counterpart, the intervals do not suggest that the difference between HiC and linear is statistically significant, except in Chromosomes 1, 9 and 21. However, the differences in these SIEs should not be quantitatively compared (e.g. measuring the difference or ratio of these SIE’s), as they all have different number of genes and connectivity. There does not seem to be correlation between the parameter estimates from the PhiMRF model and the network size or connectivity (Figure 2a, top).

**Figure 2:**
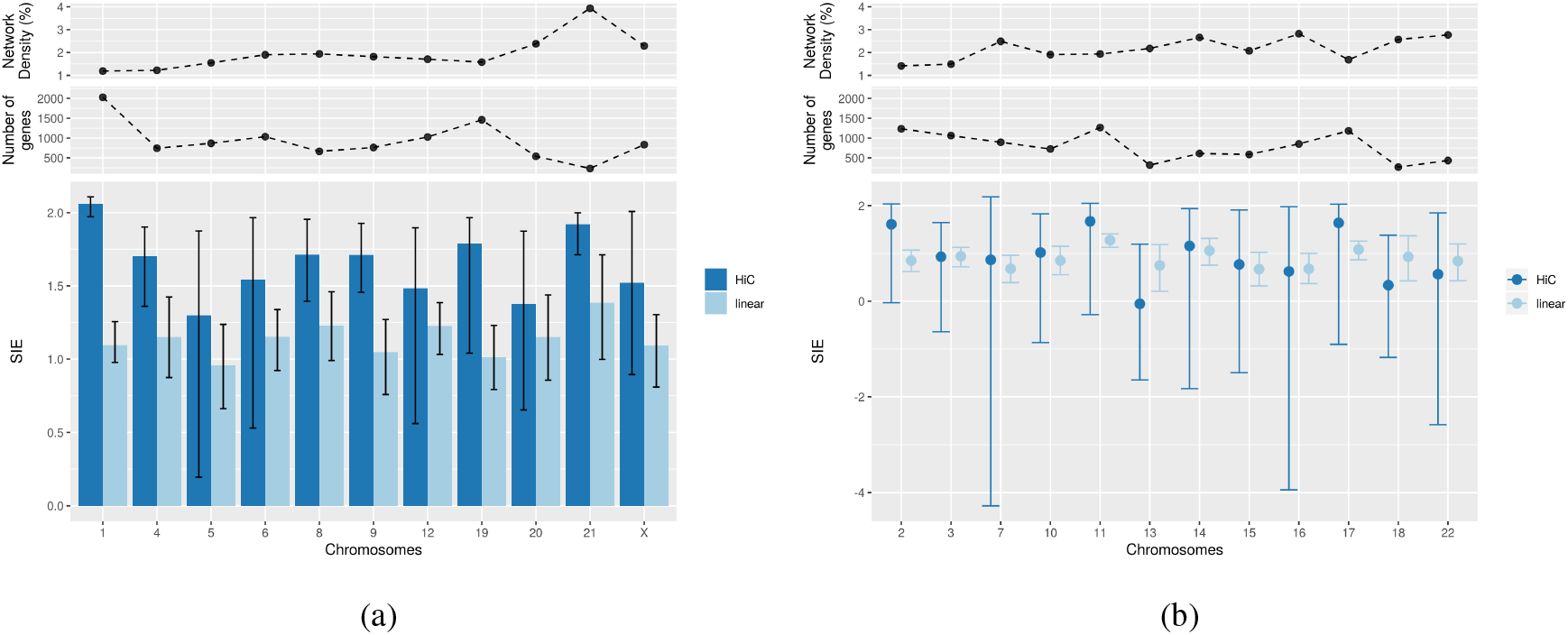
Spatial Interaction Estimate (SIE) for whole chromosomes. Top panels depicts the network properties of each chromosome. Network density is defined as the percentage of actual edges versus number of possible edges. **(a)** Chromosomes with 95% credible interval above zero. Height of bar is SIE, and error bars are the 2.5% and 97.5% quantiles from the posterior distributions. **(b)** Chromosomes with 95% credible interval including 0. Error bars are the 2.5% and 97.5% quantiles from the posterior distributions.

Twelve chromosomes did not show meaningful spatial dependency in gene expression (Figure 2b, Supplementary Tables S1 and S2). Despite only half of the chromosomes showing 3D spatial dependency, all 23 chromosomes show meaningful positive linear dependency in gene expression. In other words, the expression level of a gene is predictive of the expression levels of (at most) two other genes that are within 10k base pairs upstream or downstream from it. This is a confirmation for the efficacy of our model as it suggests that the linear dependency in gene expression is stable and detectable, in line with our existing perception of the transcription mechanism. All chromosomes have comparable estimated basal expression rates and conditional variance (Supplementary Table S1). The large estimated conditional variance is indicative of the large variation in gene expression within a chromosome.

#### Within Topologically Associating Domains

Topologically Associating Domains (TAD) are megabase-sized spatial structures in the chromosomes observed from HiC data, displaying significantly more frequent interactions within than outside these domains [18] (Figure 3a). Evidence shows that enhancer-promoter interactions are constrained within TADs [11], and genes within TADs are more active in transcription than genes in TAD boundaries [18]. However, it is unclear how TAD structures causally affect gene expression levels, especially on a global scale, instead of individual cases [30, 21]. Here we investigate the level of gene expression spatial dependency for genes located within TAD boundaries (Supplementary Table S3). We used Arrowhead[17] as our TAD caller, while several algorithms are available for the computational identification of TADs based on HiC data with varying levels of accuracy [31].

**Figure 3:**
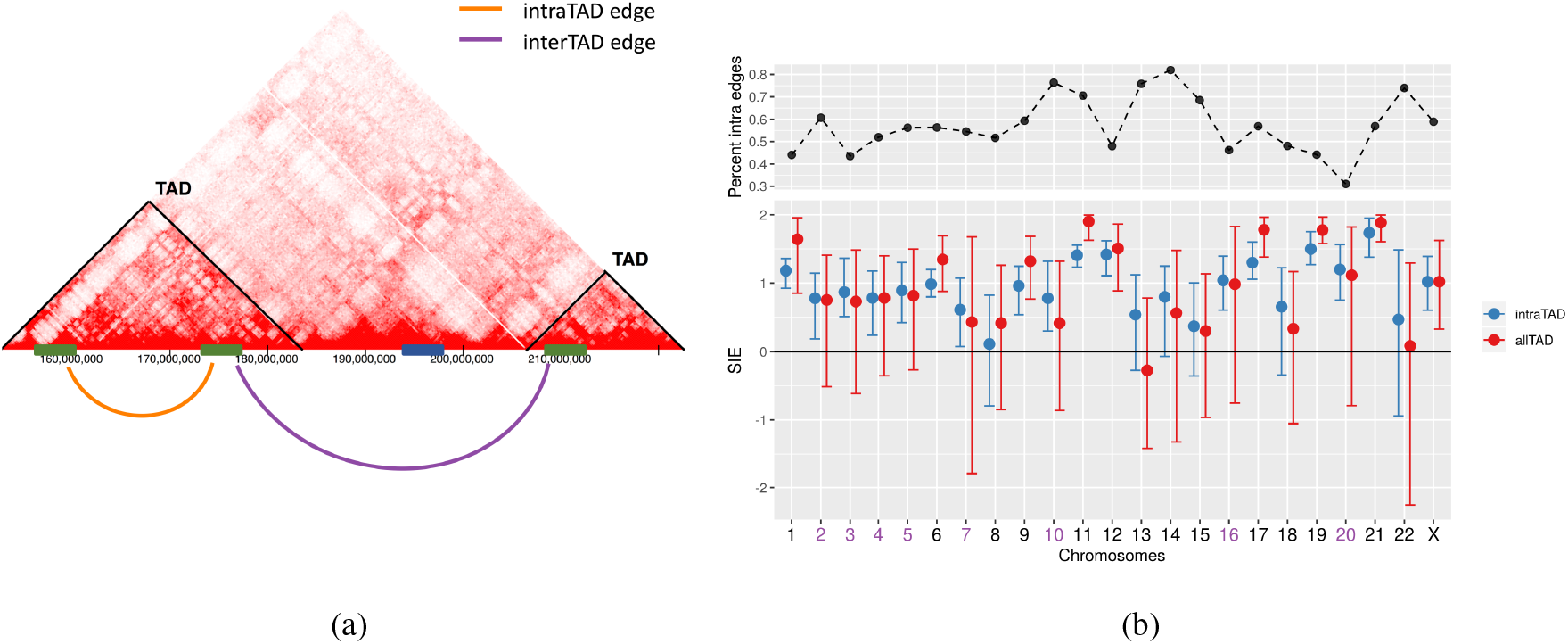
Spatial dependency in TAD gene expression. **(a)** Illustration of intra-TAD edges. **(b)** SIE of TAD genes using intraTAD versus allTAD edges. Blue: only includes edges within one TAD (intraTAD). Green: includes all edges connecting all TAD genes (allTAD).

Genes within TADs show more network clustering [32], but do not seem to exhibit extreme hub nodes versus non-TAD genes (Supplementary Figure S2). To further investigate the genes within TADs, we isolated those edges that are located within TADs as well, i.e. interactions that connect two genes within the same TAD. About half of the degree for genes located within TADs are intra-TAD edges (Figure 3b, top). Then we ran PhiMRF to detect spatial dependency of gene expression on these TAD genes, using only the edges within each individual TAD (**intra-TAD** edges), where the network is essentially made up of a group of connected components that represent each TAD. Seventeen chromosomes, with the exception of 8, 13, 14, 15, 18, 22, show meaningful spatial dependency when only including intra-TAD edges, while only nine chromosomes show positive spatial dependency when including both intra-TAD and inter-TAD edges. We also ran the model on TAD genes with only **inter-TAD** edges, to rule out the possibility that the difference in SIE is due the number of edges in the network. The dataset using inter-TAD edges display similar results as the dataset using all edges (Supplementary Figure S3). We therefore conclude that for some chromosomes, limiting the interactions to within each TAD results in a more detectable effect of spatial dependency. Intra-TAD interactions are more important than inter-TAD interactions when it comes to regulating coordinated gene expression levels. Such effects might be a result of TAD boundaries acting as insulating barriers.

To completely rule out the effect of neighborhood size (edge count), and to further validate our model, in the next step, we carried out an *in silico* experiment to disrupt the TAD boundaries by randomly sampling edges in our HiC network. In other words, we randomly permuted the order of the edges in each HiC network to create a reference distribution. For each of these 100 random samples, edges are randomly designated between any two genes. The total number of random edges is equal to the number of intra-TAD edges. We then fit PhiMRF for all 100 networks and obtain the SIE’s (Figure 4, Supplementary Figure S4). For most chromosomes, the SIE from our observed model is significantly higher than our reference distribution built from 100 randomly sampled networks. Moreover, the observed PhiMRF model often reports significantly lower remaining conditional variance when compared with randomly permuted networks. This is because some of the variance is accounted for by the spatial dependency, through the observed HiC network. At the same time, the randomly permuted networks cannot account for the spatial dependency, therefore having higher estimated variance. This serves as a validation that PhiMRF is picking up real spatial dependency signal instead of noise in the data. More importantly, the permutations could be viewed as a disruption of TAD boundaries, artificially connecting genes located in different TADs together. The PhiMRF results of such disruption demonstrated that these artificial connections could not explain the variation of gene expression. Therefore, we have proven the native organization of TADs is non-random and has significant effects in gene expression.

**Figure 4:**
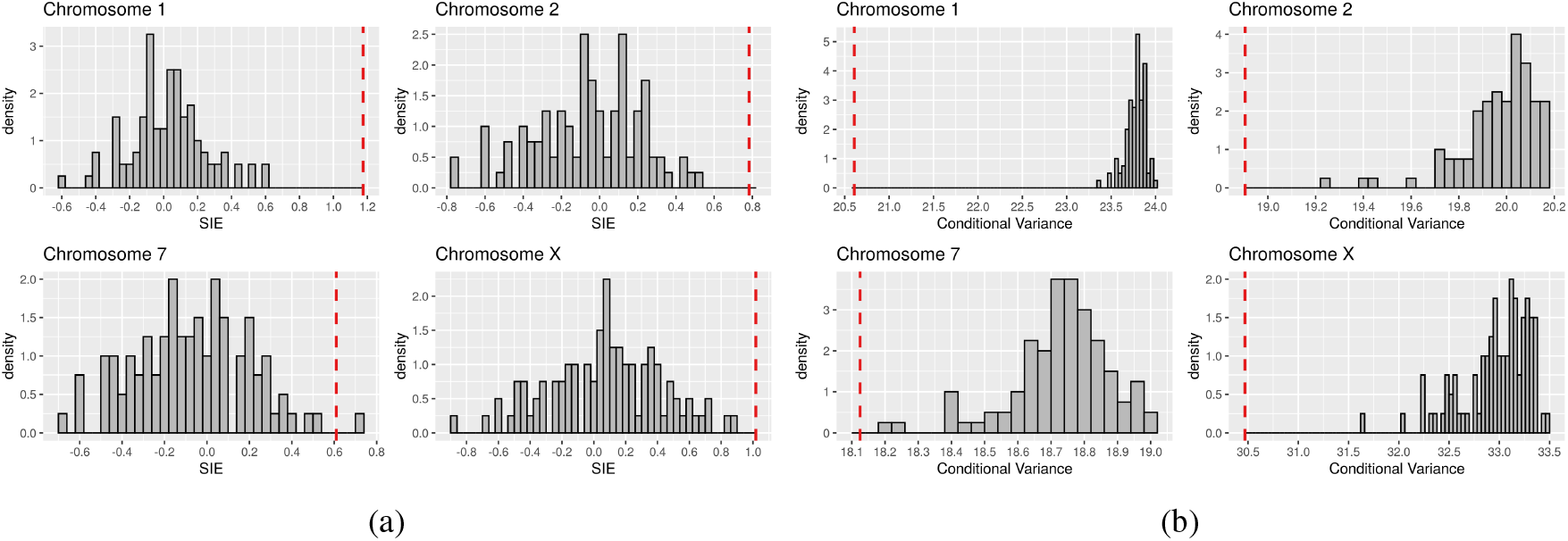
Permutation test for spatial dependency in TAD gene expression. **(a)** Histogram of SIE for 100 randomly permuted networks for all TAD genes in four chromosomes. Red dashed line is the observed SIE from the non-random HiC network. **(b)** Histogram of 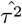 for 100 randomly permuted networks for all TAD genes in four chromosomes. Red dashed line is the observed 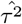 from the non-random HiC network. The permutation is carried out using an Erdős-Rényi algorithm with equal probability for any possible edge to be sampled. Total number of actual edges in each random graph is equal to the number of intra-TAD edges in the observed HiC network.

### 2.3 Inter-chromosomal dependency

Many HiC studies are focused on intra-chromosomal interactions since chromatin looping and TADs are important mechanisms for gene regulation [11, 17]. Moreover, it was initially observed that different chromosomes tend to occupy different territories in the nucleus with rare inter-chromosomal interactions [33]. Despite the discrete chromosome territories, there are about 5-10% of chromosome intermingling [34], and intermingling has a strong correlation with gene expression [35, 36]. The inter-chromosomal HiC gene networks are derived the same way as the intra-chromosomal ones, but only using inter-chromosomal HiC interactions. In other words, no two genes from the same chromosome are considered to be connected. For a total of 253 pairs of chromosomes, only 32 (12.64%) pairs do not show meaningful positive spatial dependency. In general, the lower the SIE, the larger the intervals (Figure 5, Supplementary Table S4). When spatial dependency is less detectable, it manifested in both the effect sizes and the statistical significance. The chromosome pair with the highest SIE is Chromosome 9 and Chromosome 21, followed by Chromosome 9 and Chromosome 13. Some evidence suggests that Chromosome 9 is located in the center of the nucleus [37], which may explain the high inter-chromosomal spatial dependency of its gene expression. Another chromosome that appeared twice in the top ten list is chromosome 19, which has also been shown to locate in the center of the nucleus [38]. Chromosome 1 is the only chromosome that appeared three times in the top ten list, which might be attributed to its exceptional length and gene count. We then incorporated all inter-chromosomal and intra-chromosomal HiC edges to all 19631 genes in the genome. Among the total 2,368,756 edges, 176,282 (7.44%)are intra-chromosomal edges. The global gene network has a SIE of 2.561(2.551, 2.570), confirming spatial dependency of gene expression as a global effect, observed in the whole genome. The global dataset has a basal expression rate of 3.039 (2.909, 3.176) and conditional variance of 22.337 (21.607, 23.227).

**Figure 5:**
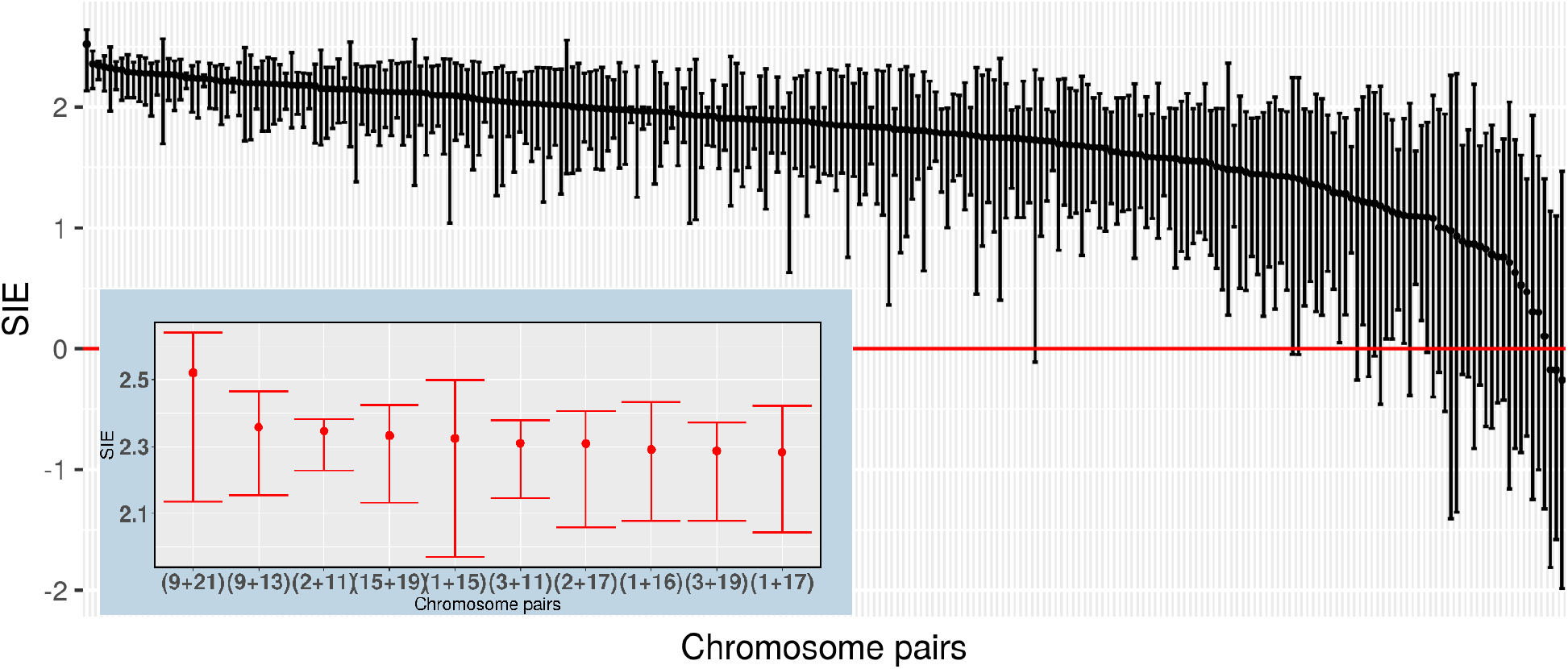
SIE of chromosome pairs using only inter-chromosomal HiC interactions. For each pair of chromosomes, PhiMRF ran on a dataset where all genes in both chromosomes are included, while only inter-chromosomal edges are included. From left to right, chromosome pairs ranked by highest SIE to lowest. **Background:** SIE and 95% credible interval of all 253 chromosome pairs. **Zoomed overlay:** SIE and 95% credible interval of the top ten chromosome pairs with the highest SIE.

### 2.4 Functional gene groups

Next we tested the hypothesis that spatial dependency of gene expression is correlated with the function of the genes. Evidence suggest that co-functioning genes tend to cluster in space, and function is closely correlated with expression levels, since co-functioning genes sometimes need to be co-transcribed [39, 22].

Having developed the PhiMRF framework to understand the relationship between expression and spatial organization, we can apply it to any group of genes, to see whether spatial dependency is particularly strong for certain functions, pathways or phenotypes. The implication of a large SIE for a functional group is that the 3D organization of the genome plays a role in the regulation of such function. Here we demonstrate this application of PhiMRF using the Gene Ontology (GO) consortium as the controlled vocabulary for functional annotation.

We picked the top fifteen GO terms by count in the Biological Process aspect of GO (BPO) and collected all the genes associated with each GO term (Supplementary Table S5). These terms include functions like transcription regulation, protein phosphorylation etc. The size of the gene groups varies from 504 genes to 167 genes.

Groups of genes associated with positive/ negative regulation of transcription by RNA polymerase II (GO:0045944, GO:0000122),G protein-coupled receptor signaling pathway (GO:0007186), neutrophil degranulation (GO:0043312), and negative regulation of cell population proliferation (GO:0008285) shows statistically meaningful spatial dependency (Figure 6, Supplementary Table S6).

**Figure 6:**
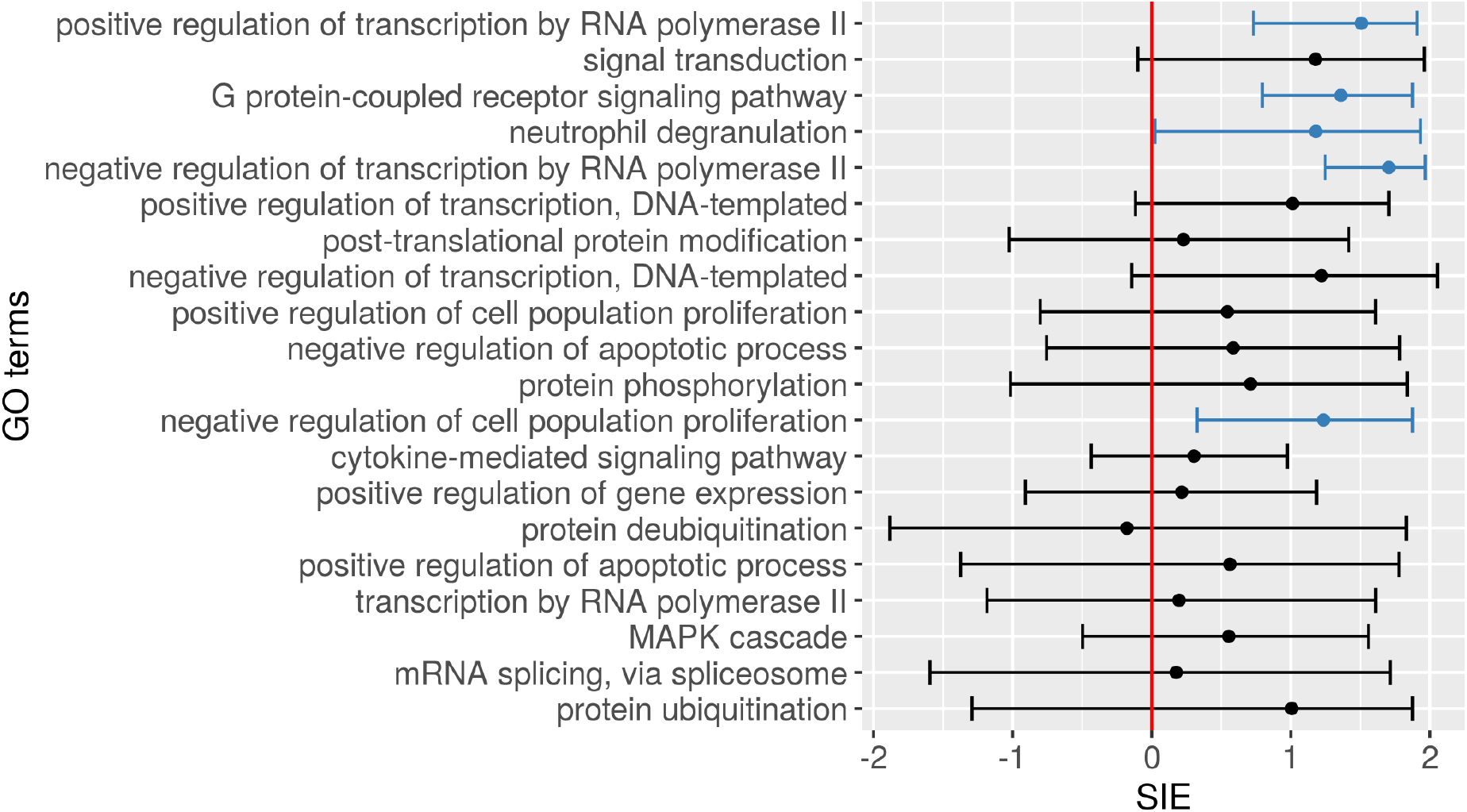
SIE of functional gene groups. SIEs and 95% credible intervals obtained for each group of genes associated with the top 20 GO BPO terms. For each group of genes, both intra-chromosomal and inter-chromosomal HiC interactions are included. The GO terms are ranked by the number of genes the annotate in decreasing order from top to bottom. Meaningful spatial dependencies are marked in blue.

The interpretation of these positive results is the interplay of the three important factors of a gene: its spatial location, its expression level and its function. A positive SIE for a functional gene group means that genes that are part of this particular biological process have co-dependent expression levels based on their spatial locations. In order for these genes to carry out the same function together, regulation of the expression of these genes takes advantage of the 3D genome structure.

### 2.5 Cell line difference

We replicated intra-chromosomal and inter-chromosomal PhiMRF analyses in another cell line, GM12878. From Differential Expression (DE) analysis using DESeq2 [27], the two cell lines shows different expression landscape (Supplementary Figure S6). About half of the genes in each chromosome are differentially expressed (Supplementary Table S7), summing to a total of 10,812 DE genes. The HiC network structures for the two cell lines are also different. We acquired the *combined* HiC contact matrices containing both primary and replicate HiC experiments. Under the same edge inference criteria (Methods), the GM12878 HiC networks contains about twice more edges than the IMR90 networks (Supplementary Table S7). Edges in GM12878 covers all of the edges in IMR90 while adding a lot more others, giving the networks much higher density (Supplementary Figures S7a and S7b, top).

Despite the differential expression and different network structures, we observe similar patterns of spatial dependency in the GM12878 cell line. Eight chromosomes: 1, 4, 5, 11, 14, 15, 21 and X show meaningful spatial dependency, with significant larger SIE’s than their linear counterparts (Supplementary Figure S7a).

Chromosomes 1, 4, 5, 21 and X showed meaningful spatial dependency in both cell lines, indicating that this regulation mechanism may be common and essential among all cell lines and tissue types. These five chromosomes are not different from other chromosomes in terms of gene count, edge ratio or number of DE genes. More experiments are need to determine if there is anything special about these five chromosomes. For those chromosomes that did not show meaningful intra-chromosomal spatial dependency in GM12878, they exhibited large credible intervals, due to the large number of edges, making the dependencies less concentrated and hence harder to detect. In both cell lines, the same structure underlines the linear gene networks. Both cell lines exhibit meaningful and consistent linear spatial dependency across all chromosomes, suggesting that the mechanism of coordinated expression in linear neighboring genes is still preserved even when the expression levels are different for these genes.

In terms of inter-chromosomal dependencies, only 60 (23%) out of the 253 chromosome pairs showed spatial dependency in GM12878, significantly less (*p <* 0.01, test of proportions, two-sided) than that of IMR90 (Supplementary Figure S8). However, the top ten most significant pairs reveals some familiar chromosomes. For example, chromosome 19 appeared three times in the top ten list of most spatially dependent chromosome pairs in GM12878, as was the case in IMR90.

## 3 Discussion

We have developed a novel probabilistic model for gene expression incorporating spatial dependency. By applying this model to RNA-seq and HiC data, we get our first peek into how the transcriptome is shaped by the 3D location of the genes. The different expression levels of genes can be partially explained by the spatial location of the genes. The quantitative measurement of such effects provides unprecedented insight into the relationship and interplay among chromosomal organization, gene expression and functionality.

### Insulating effects of TADs

It has been shown that TADs play a role in the regulation of gene expression [18]. For example, the expression profiles of genes whose promoters are located within the same TAD are more correlated during cell differentiation than those of genes not in the same TAD [36]. One hypothesis is the regulation is weakened by the insulating effects of TAD boundaries. TAD boundaries are enriched with CTCF proteins and CTCF binding sites, which are known to help shape the structure of the genome [18]. Our study confirms that the expression of intra-TAD genes shows spatial dependency on a global scale, strengthening the hypothesis that TADs are an integral part of the gene expression regulation mechanism. Interestingly, since we are able to examine the 3D spatial interactions within each TAD, we discovered that for eight chromosomes, spatial dependency is only detectable when the interactions are confined to TADs (Figure 3b). If we include only inter-TAD edges (interactions between two genes on two different TADs), we observe meaningful spatial dependency only in five chromosomes, compared with seventeen chromosomes for intra-TAD edges (Supplementary Figure S3). All edges present in the HiC gene network represent spatial proximity of these genes in the 3D space. However, if some of these edges are insulated or blocked by boundary elements such as CTCF proteins, then it makes sense that the spatial dependency of gene expression is no longer detectable when we include these edges. Even though genes connected by these inter-TAD edges are still spatially close, their expression are no longer coordinated due to the insulating effect of TAD boundaries.

### High inter-chromosomal dependency

The rapid pace of development of Chromosome Conformation Capture (3C) technology enables us to obtain detailed understanding of intra-chromosomal architecture that shapes the regulatory landscape of the genome, but inter-chromosomal interactions and their functions are studied less and still poorly understood [40]. Many known inter-chromosomal interactions are not detected by HiC experiments, although they can be as stable as intra-chromosomal contacts [41]. Maass *et al* reasoned that the lack of significant detection of the inter-chromosomal interactions is due to the different distance scale of the inter– versus the intra-chromosomal ones [41]. We acknowledge the bias that the HiC technique has against inter-chromosomal interactions. To mitigate this bias, we used a high resolution (10kb) HiC interaction map, together with the soft thresholding of HiC interactions. The high resolution map enabled us to pool information from several loci for one gene, while the soft thresholding allowed us to infer a high interaction frequency based on individual chromosome pairs and not any absolute distance across the genome. However, even with such measures, network density is still lower for gene networks of inter-chromosomal pairs than for intra-chromosomal gene networks (Supplementary Figure S5), demonstrating the need for caution when using and interpreting HiC data. Despite the relative low network density for inter-chromosomal HiC gene networks, we were able to observe high inter-chromosomal spatial dependency in gene expression for the majority (87.35%) of chromosomal pairs in IMR90 cells (Figure 5). These results suggest extremely long-range and intra-chromosomal gene interactions on different chromosomes as a commonly occurring regulation mechanism for gene expression.

In summary, we have developed a hierarchical Markov random field (PhiMRF) model to explain the variation in gene expression. PhiMRF can be further applied to gene groups that are functionally enriched, genes in the same biological pathway, genes that are causal for a certain disease or phenotype, and so on. In doing so, we are essentially looking at the regulation mechanism for such gene groups to perform a function or set of functions. A meaningful spatial dependency would indicate that the regulation of such function or disease involves the 3D genome architecture as one of the regulation mechanisms, allowing biologists to explore new directions when studying the functions, pathways or diseases of interest.

## 4 Methods

The gene expression data used in this study comes from the ENCODE project (https://www.encodeproject.org) with the following identifiers ENCFF353SBP and ENCFF496RIW for IMR90 and ENCFF680ZFZ and ENCFF781YWT for GM12878. These are each two biological replicates of total RNA-seq experiments on the IMR90 and GM12878 cell lines. Genes are mapped to Ensembl stable IDs with coordinates from Ensembl release 90, which uses the GRCh38.p10 human genome assembly.

### 4.1 Bayesian Inference

The goal of our Bayesian framework is to simulate from the posterior distributions *p*(*α*|***y***), *p*(*η*|***y***) and *p*(*τ*^2^|***y***). The properties of these distributions directly answer the biological questions from which we abstracted the stochastic model. The overall strategy is to simulate from the joint posterior distribution of *p*(*α, η, τ*^2^, ***w**|**y***) using a Gibbs sampler, where we sequentially simulate from each of the full conditional posteriors of our parameters as follows.

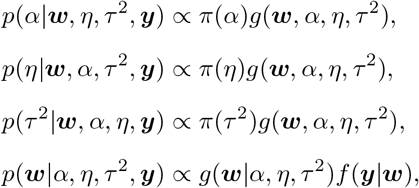

where *π*(*α*), *π*(*η*) and *π*(*τ*^2^) are Uniform prior distributions, *f* (***y**|**w***) is the Poisson distribution for observed data ***y***, and *g*(***w**|α, η, τ*^2^) is the marginal distribution for ***w***. This marginal distribution is not readily available from our model specification, since ***w*** is only conditionally specified. However, it can be constructed from the conditional distributions using a negpotential function [42] (Supplementary Methods). Moreover, the negpotential introduces an intractable constant to the posterior that cannot be dropped, so we use the *double* Metropolis-Hastings algorithm to simulate from these posteriors (Supplementary Methods).

#### Prior distributions

The prior used for *α* is a Uniform distribution *U* (*−*10, 10). The prior used for *η* is a Uniform distribution over the parameter space of *η*. The parameter space of *η* is directly calculated from the neighborhood adjacency matrix, as the inverse maximum and minimum of its eigenvalues (Supplementary Methods). We assume that *τ* follows a prior uniform distribution *U* (0, 10) and derive the prior distribution for *τ*^2^ in the model as 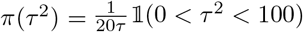.

#### Iterations

For each intra-chromosomal (including the linear baseline) and pairwise inter-chromosomal HiC dataset, we ran the double Metropolis-Hastings algorithm through 5000 iterations, with 1000 burn-in iterations. All TAD related datasets, the whole-genome dataset and the functional datasets went through 2000 iterations with 400 burn-in. The variances for the jump proposal distributions (Supplementary Methods) were chosen through multiple rounds of initial testing to ensure that the jump frequencies fall within 15% to 40% for randomly selected datasets in each group of datasets, and that the MCMC simulations converge within the number of iterations used.

### 4.2 HiC Data processing

#### Normalization

Raw observed HiC data for the IMR90 and GM12878 cell from Rao *et al*[17] with 10kb resolution were used. The KR normalization technique was applied. The goal of the normalization is to remove one-dimensional bias in HiC counts. On Chromosome 9 of IMR90 at this resolution, the KR algorithm did not converge on that particular matrix, this is likely due to sparsity of the matrix. In this case raw counts are used. Interactions between the same locus are eliminated to avoid bias towards neighboring genes.

#### Gene Mapping

Interactions are observed for every consecutive 10k bins (loci) on the entire human genome (except chromosome Y), while expression are generally considered for genes located intermittently on the genome. Since each bin is 10kb in size, one gene is often mapped to multiple bins. The interaction between two genes is then decided by all of the interacting bins that overlap with the pair of genes. An overlap is when a bin shares more than 10% of base pairs with a gene (Supplementary Figure S1). For example, if gene 1 overlaps with bin A and bin B, while gene 2 overlaps with bins C, D and E, then the interaction between gene 1 and gene 2 are considered as the pool of interactions A-C, A-D, A-E, B-C, B-D and B-E. The goal of such gene-bin mapping is to inform a gene network where edges connect gene pairs that are close in 3D space while eliminating potential bias from individual loci. We found that most gene pairs overlap with a pool of less than five bin pairs, for both intra-chromosomal and inter-chromosomal gene pairs. Therefore, we adopted simple metrics to summarize these pools of interactions instead of using more complicated parametrizations (e.g. a t test). Four metrics are considered, mean, median, max and min to summarize this pool of interactions. The computed metric for each pair of genes is then compared to a threshold to decide whether two genes are neighbors.

### Soft threshold

The mean (median, max or min) of the pool of interactions for a gene pair is compared with a threshold to determine whether there is an edge between the gene pair. The higher the interaction score, the closer in proximity. The threshold is determined as a 90% quantile of *all* locus-locus interactions found in that chromosome or chromosome pair. Therefore, the threshold changes from chromosome to chromosome, eliminating chromosomal bias. Out of the four metrics, the min metric is the most conservative, as it requires that all interactions in the pool be larger than the threshold, to consider the gene pair to be neighbors, while the max metric only requires that one interaction out of the pool be larger than the threshold. Mean is our main metric, and all subsequent studies presented in the main text uses the mean metric. Intra-chromosomal datasets are repeated using the other three metrics (median, min and max) as well and presented in Supplementary Tables S9, S10 and S11.

### 4.3 Functional Annotations

We used the UniProt Gene Ontology Annotation (downloaded on September 16, 2019) to extract all Gene Ontology BPO terms annotating human gene products. We only extracted the annotations with experimental evidence, with the following evidence codes: EXP, IDA, IPI, IMP, IGI, IEP, TAS and IC. When counting the most annotated terms, we decided to *not* propagate the annotations through the hierarchical structure of the ontology. An example of a propagation is where the ontology structure specifies that *carbohydrate phosphorylation* (GO:0046835) "is a" *phosphorylation* (GO:0016310). If a protein is annotated with the former, a more informative term, then it is automatically annotated with the latter, a less informative term. Since we are ranking the GO terms by the number of proteins they annotate, if the annotations are propagated, the top ones will be the very general functions like "cellular process", that are in general not of interest to researchers (Supplementary Table S5). Therefore, we have elected to rank the GO terms by the number of proteins they directly annotate. These leaf annotations are directly curated from published experiments, which is evidence that these terms are of interest to some researchers.

### 4.4 Data and software availability

The source code, prerequisites and installation guide, as well as a Docker image for the R package PhiMRF are available at https://github.com/ashleyzhou972/PhiMRF under GPL-2 license.

The scripts for data processing are available at https://github.com/ashleyzhou972/bioMRF under a GPL-2 license.

Full intermediate and final results are available at https://doi.org/10.6084/m9.figshare.11357321.v4.

