## Supplementary file for "Hierarchical Markov Random Field model captures spatial dependency in gene expression, demonstrating regulation via the 3D genome"

S2 **Connectivity measurements for genes within and outside of TADs. Top:** Max degree is the maximum number of edges for a node in the network. **Middle:** Network density is defined as the percentage of actual edges versus number of possible edges. **Bottom:** Clustering Coefficient (also called transitivity) is the ratio between number of closed triplets in the network and number of all triplets. A triplet is three nodes that are connected by either two (open) or three (closed) edges. . . . . . 5
S3 **SIE of TAD genes using intraTAD versus interTAD edges.** Blue: only includes edges within one TAD (intraTAD). Green: only includes edges connecting genes in two different TADs genes (interTAD). . . . . . 6
S4 **Histogram of $\hat{\alpha}$ for 100 randomly permuted networks for all TAD genes in four chromosomes.** Red dashed line is the observed $\hat{\alpha}$ from the non-random HiC network. . . . . 7

S5 **Histogram of network properties for 23 intra-chromosomal gene networks versus 253 inter-chromosomal gene networks. Left:** Network density. **Right:** Edge to Node ratio . . . . . . . . . . . . . . . . . . . . . . . . . . . . . . . . . . . . . . . . . . . . 8
S6 **Differential Expression analysis on GM12878 and IMR90 cell lines.** DE analysis was done using the DESeq2[1] package.Red: genes that are differentially expressed. Green: genes that are not differentially expressed. . . . . . . . . . . . . . . . . . . . . . . . . . . 9
S7 **GM12878: Spatial Interaction Estimate (SIE) for whole chromosomes.** Top panels depicts the network properties of each chromosome. Network density is defined as the percentage of actual edges versus number of possible edges. **(a)** Chromosomes with 95% credible interval above zero. Height of bar is SIE, and error bars are the 2.5% and 97.5% quantiles from the posterior distributions. **(b)** Chromosomes with 95% credible interval including 0. Error bars are the 2.5% and 97.5% quantiles from the posterior distributions. . . . . . . . . . 10
S8 **GM12878: SIE of chromosome pairs using only inter-chromosomal HiC interactions.** For each pair of chromosomes, PhiMRF ran on a dataset where all genes in both chromosomes are included, while only inter-chromosomal edges are included. From left to right, chromosome pairs ranked by highest SIE to lowest. **Background:** SIE and 95% credible interval of all 253 chromosome pairs. **Zoomed overlay:** SIE and 95% credible interval of the top ten chromosome pairs with the highest SIE. . . . . . . . . . . . . . . . . . . . . . . 11

S10 **Posterior parameter estimates using the min metric**. For each chromosome, when summarizing the pool of interactions for each gene pair, **min** is used as the summary statistic. 26
S11 **Posterior parameter estimates using the max metric**. For each chromosome, when summarizing the pool of interactions for each gene pair, **max** is used as the summary statistic. 27

### Supplementary Methods

#### Conditionally specified models

What sets Equation (1) apart from the Gaussian assumption for a simple linear regression is that the distribution for $w_i$ is conditionally specified, as opposed to the marginally specified model, e.g. $Y_i \sim N(\mu_i, \tau^2)$. Moreover, in a simple linear regression, the next step of the model is to regress the Gaussian mean against independent variable $X$ (e.g. $\mu_i = X_i\beta$), while in our *autoregressive* model, the Gaussian mean is regressed against its immediate neighbors $\boldsymbol{w}(N_i)$, shown in Equation (2). The parameter $\eta$ in Equation (2) is analogous to the regression coefficient $\beta$ in a simple linear regression, that symbolizes effect of the spatial autocorrelation. $\frac{1}{|N_i|+|N_j|}$ is a normalizing constant that adjusts the effect of different number of neighbors for different gene $i$'s. $\alpha$ is a parameter analogous to the intercept in a simple linear regression model and $\tau^2$ is analogous to the variance parameter $\sigma^2$ in a simple linear regression.

#### Constructing the marginal distributions

We can derive the full conditional posterior distribution of $\alpha$ as $p(\alpha|\eta, \tau^2, \boldsymbol{y}, \boldsymbol{w}) \propto \pi(\alpha)g(\boldsymbol{w}|\alpha, \eta, \tau^2)$, where $\pi(\alpha)$ is the prior distribution for $\alpha$. Note that this calls for the *marginal* distribution of $\boldsymbol{w}$ ($g(\boldsymbol{w}|\alpha, \eta, \tau^2)$). Since our model is conditionally specified, this distribution is not readily available. For certain MRF models, the marginal distributions can be constructed from conditionals using the negpotential function [2] as follows.

$$
\begin{aligned}
&p(\alpha|\eta, \tau^2, \boldsymbol{y}, \boldsymbol{w}) \\
&\propto \pi(\alpha)g(\boldsymbol{w}|\alpha, \eta, \tau^2) && (1)\\
&= \pi(\alpha)\frac{exp(Q(\boldsymbol{w}|\alpha, \eta, \tau^2))}{\int exp(Q(\boldsymbol{w}|\alpha, \eta, \tau^2))d\boldsymbol{w}} && (2)\\
&= \pi(\alpha)C(\alpha)exp(Q(\boldsymbol{w}|\alpha, \eta, \tau^2)) && (3)
\end{aligned}
$$

where $Q(\boldsymbol{w}|\alpha, \eta, \tau^2))$ is the negpotential function as defined in [2] and can be constructed using $Q(\boldsymbol{y}|\boldsymbol{\theta}) = \sum_{1\leq i\leq j} H_i[y(s_i)|\boldsymbol{\theta}] + \sum\sum_{1\leq i<j\leq n} H_{i,j}[y(s_i), y(s_j)|\boldsymbol{\theta}]$ and $C(\alpha) = 1/\int exp(Q(\boldsymbol{w}|\alpha, \eta, \tau^2))d\boldsymbol{w}$ is an intractable constant. Since our posterior distribution for $\alpha$ (and likewise for $\eta$ and $\tau^2$) contains an intractable constant that cannot be dropped, we utilized the double Metropolis-Hastings algorithm[3] that allows us to sample from the posterior distribution using an MCMC procedure different from a regular Metropolis-Hastings algorithm.

#### Double Metropolis-Hastings algorithm

Here we outline the steps of a double MH algorithm using the full conditional posterior distribution of $\alpha$ (equation 3).

Let $t$ denote the index of MCMC iteration, and let $\alpha_t$ denote the current state of the Markov chain.

**Step 1.** Simulate a new $\alpha'$ from $\pi(\alpha)$ using the MH algorithm starting with $\alpha_t$. Note that this is a one-step MH process, instead of simulating directly from the prior distribution $\pi(\alpha)$. From a traditional MH point of view, this means treating $\pi(\alpha)$ as the invariant (target) distribution and calculation of the acceptance probability is needed.

**Step 2.** Generate an auxiliary variable $\boldsymbol{y} \sim P^{(m)}_{\alpha'}(\boldsymbol{y}|\boldsymbol{w})$, where $P^{(m)}_{\alpha'}(\boldsymbol{y}|\boldsymbol{w})$ is a MH kernel of $m$ steps. This means using the unnormalized distribution $Q(\cdot|\alpha')$ as the invariant distribution in each step of this MH process, starting with $\boldsymbol{w}$. Usually $m$ can equal to the size of $\boldsymbol{w}$, which means this step is one single round of Gibbs sampler.

**Step 3.** We accept $\alpha^{'}$, i.e. $\alpha_{t+1} = \alpha^{'}$ if the auxiliary variable is accepted, with acceptance probability $\min\{1, r(\alpha_t, \alpha^{'}, \boldsymbol{y}|\boldsymbol{w})\}$, where

$$r(\alpha_t, \alpha^{'}, \boldsymbol{y}|\boldsymbol{w}) = \frac{Q(\boldsymbol{y}|\alpha_t)Q(\boldsymbol{w}|\alpha^{'})}{Q(\boldsymbol{w}|\alpha_t)Q(\boldsymbol{y}|\alpha^{'})}.$$

The double MH algorithm can be used to simulate from the posterior distribution of a single parameter, or as employed here, to sample from conditional posteriors in an overall Gibbs sampler to sequentially sample from a joint posterior distribution.

#### Prior distributions

The prior used for $\eta$ is a Uniform distribution over the parameter space of $\eta$. The parameter space of $\eta$ is directly calculated from the neighborhood adjacency matrix, as the inverse maximum and minimum of its eigenvalues[4]. This is a very important caution in our model, where the parameter space of $\eta$ is in fact dictated by the graph structure and is not constant over different graphs. However, for most intra-chromosomal and pairwise inter-chromosomal networks, the $\eta$ parameter spaces are quite similar, with maximum at around 2.5, making the $\hat{\eta}$ values for these datasets qualitatively comparable, e.g. the level of spatial dependency in gene expression for Chromosome 1 is larger than that of Chromosome 4. However, the two $\hat{\eta}$'s from the two chromosomes are not quantitatively comparable, i.e. it is inappropriate to say the the level of spatial dependency for Chromosome 1 is 1.2 times of Chromosome 4 (Table S1), since the parameter space of the $\eta$'s are not exactly the same, hence the two numbers are not exactly on the same scale.

#### Iterations

In the initial step of a Metropolis-Hastings algorithm, random walk proposals were used for $\alpha$ and $\tau^2$, while an independence proposal was used for $\eta$. The random walk proposals enable the tuning of jump frequencies of the MCMC process by changing the variance of the random walk. In special cases such as a small number of genes with hyper connectivity, if $\alpha$ or $\tau^2$ do not converge within a large number of iterations, then the hierarchical model is considered not suitable for such dataset. No cases reported in Section 2.2 and 2.3 fall under this category.

#### Simulations

To ensure that our model can correctly represent data provided, we performed extensive simulation studies to make sure that for data generated using given parameters, our model can estimate those parameters correctly. Table S8 shows parameter estimates for data simulated with give parameters. Our model was able to recover the original simulated parameters in various parameter settings.

### Supplementary Discussion

#### Application to genomes and gene groups

HiC data in this model are represented as a spatial network, where two genes are connected if they are close in physical space. Even without HiC data, as long as any distance between each pair of genes could be represented in a network, our model can still be applied. The distance used in PhiMRF does not necessarily have to be from 3D physical space, but could be from any other types of quantifiable pairwise gene-gene interactions. The interpretation of such results depends on the specific type of gene-gene interaction used as the underlying network.

#### Co-expression and extending PhiMRF

It is important to note that PhiMRF is not modelling co-expression, since it does not account for the variability in gene expression that comes from multiple experimental conditions or samples. PhiMRF is an autoregressive model that could take one or more observations at each location. Multiple observations at one location are considered conditionally independent of each other, while co-expression (and differential expression) analyses focus on the probability space for one gene among multiple samples. Although what we find can be described as genes having *coordinated* expression based on spatial proximity, the term "co-expression" is not applicable at this stage. However, PhiMRF can be useful in co-expression (differential expression) analyses by accounting for the spatial effect in the between-sample variation.

PhiMRF also does not account for other effects that may be at play in gene expression. The basal expression level parameter $\alpha$, and the variance parameter $\tau^2$ are assumed to be the same for all genes in the genome. This assumption is crude and does not reflect the nature of the transcriptome. For example, assigning a different $\alpha$ parameter to housekeeping and non-housekeeping genes could potentially greatly reduce variation in the model since those two groups are known to have very different expression levels. Our argument for the simple model specification is that it is hard to separate these other effects from the spatial effects. At the same time, the spatial effect (close proximity of certain genes) might as well be the result of some other molecular mechanisms that also modify the gene expression profile, e.g. enhancer-promoter interactions at hub-enhancers[5]. It is not our goal to isolate the spatial effect from the other effects, rather we use the spatial effect as a proxy for other mechanisms that might be at play.

#### Producing valid posteriors

It is challenging to reproduce MCMC based results as doing so involves specific tuning of an algorithm. Our model is also restrictive in the parameter space of $\eta$, where it is dictated by the adjacency matrix structure and not a fixed value. PhiMRF should not be used on small datasets. ($< 100$ nodes). It is always good practice to check the convergence of the MCMC, and tune the parameters so that the jump rates of the MCMC fall within a reasonable range.

Figure S1: **Mapping genes to HiC observed bins.** Genes are mapped to multiple bins (loci) that have at least 10% overlap with them.

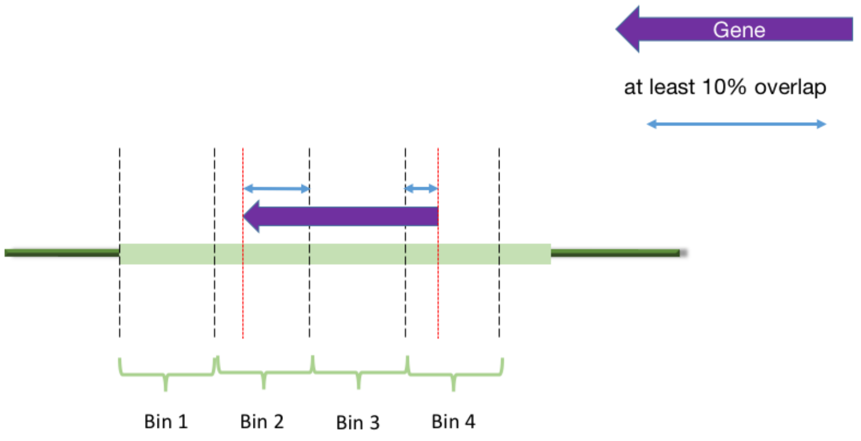

Figure S2: **Connectivity measurements for genes within and outside of TADs. Top:** Max degree is the maximum number of edges for a node in the network. **Middle:** Network density is defined as the percentage of actual edges versus number of possible edges. **Bottom:** Clustering Coefficient (also called transitivity) is the ratio between number of closed triplets in the network and number of all triplets. A triplet is three nodes that are connected by either two (open) or three (closed) edges.

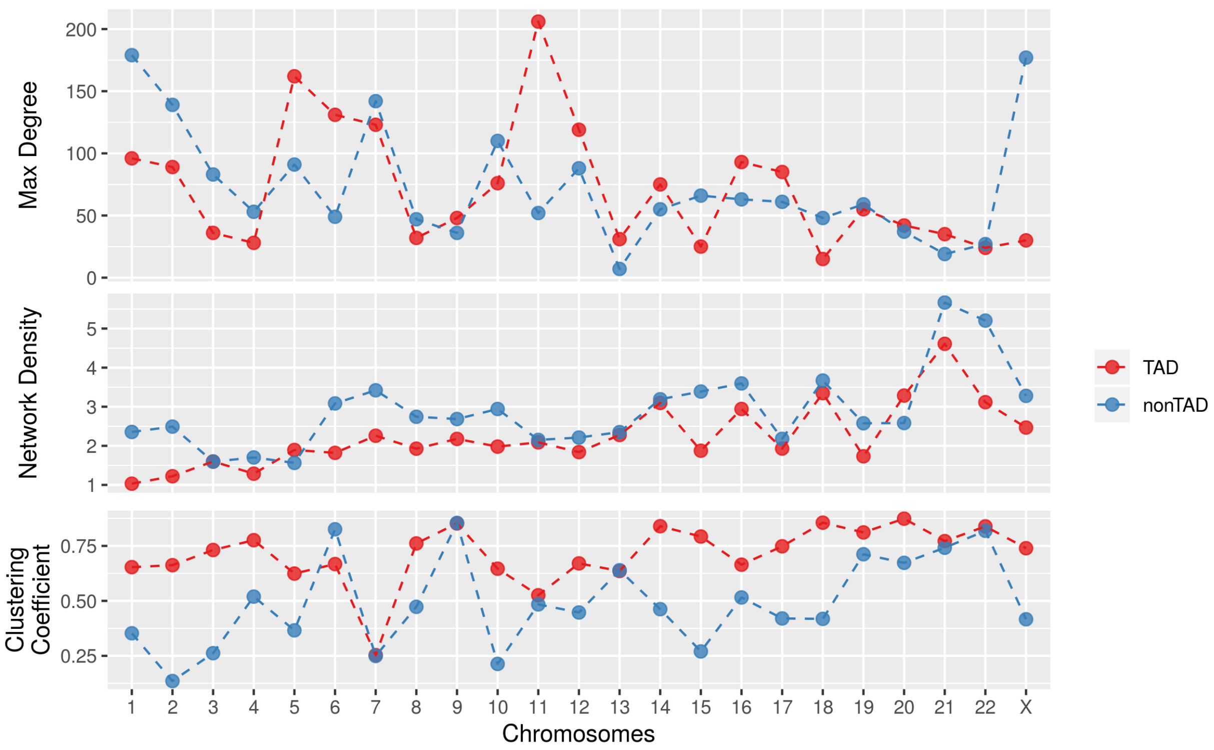

Figure S3: **SIE of TAD genes using intraTAD versus interTAD edges.** Blue: only includes edges within one TAD (intraTAD). Green: only includes edges connecting genes in two different TADs genes (interTAD).

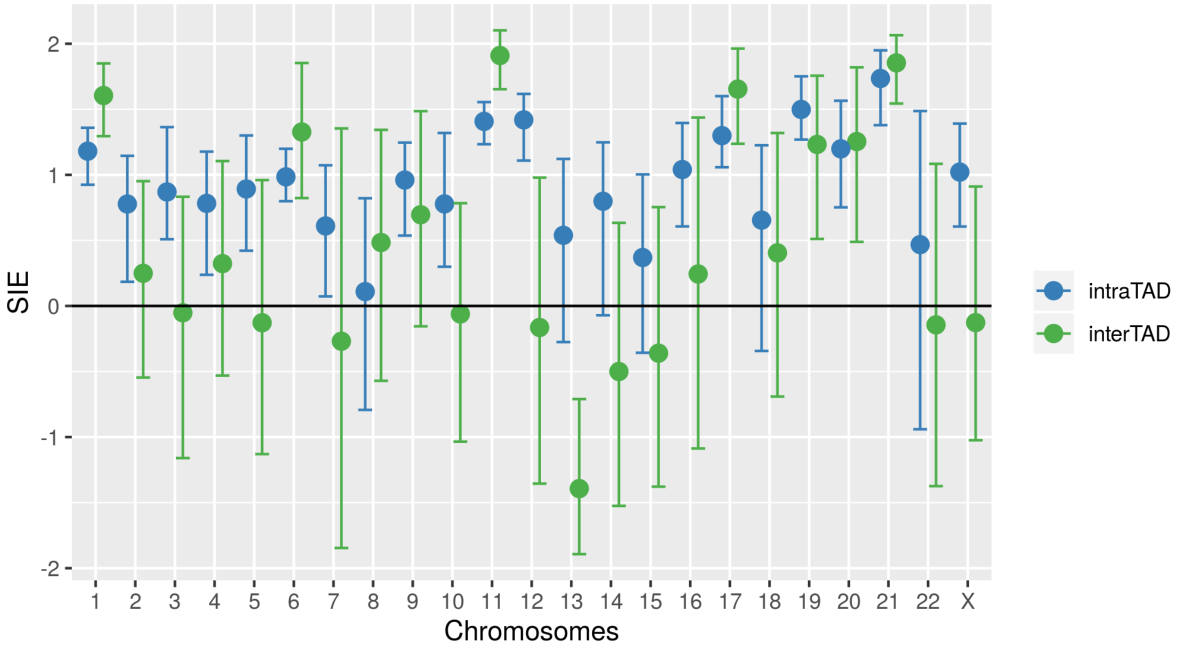

Figure S4: **Histogram of $\hat{\alpha}$ for 100 randomly permuted networks for all TAD genes in four chromosomes.** Red dashed line is the observed $\hat{\alpha}$ from the non-random HiC network.

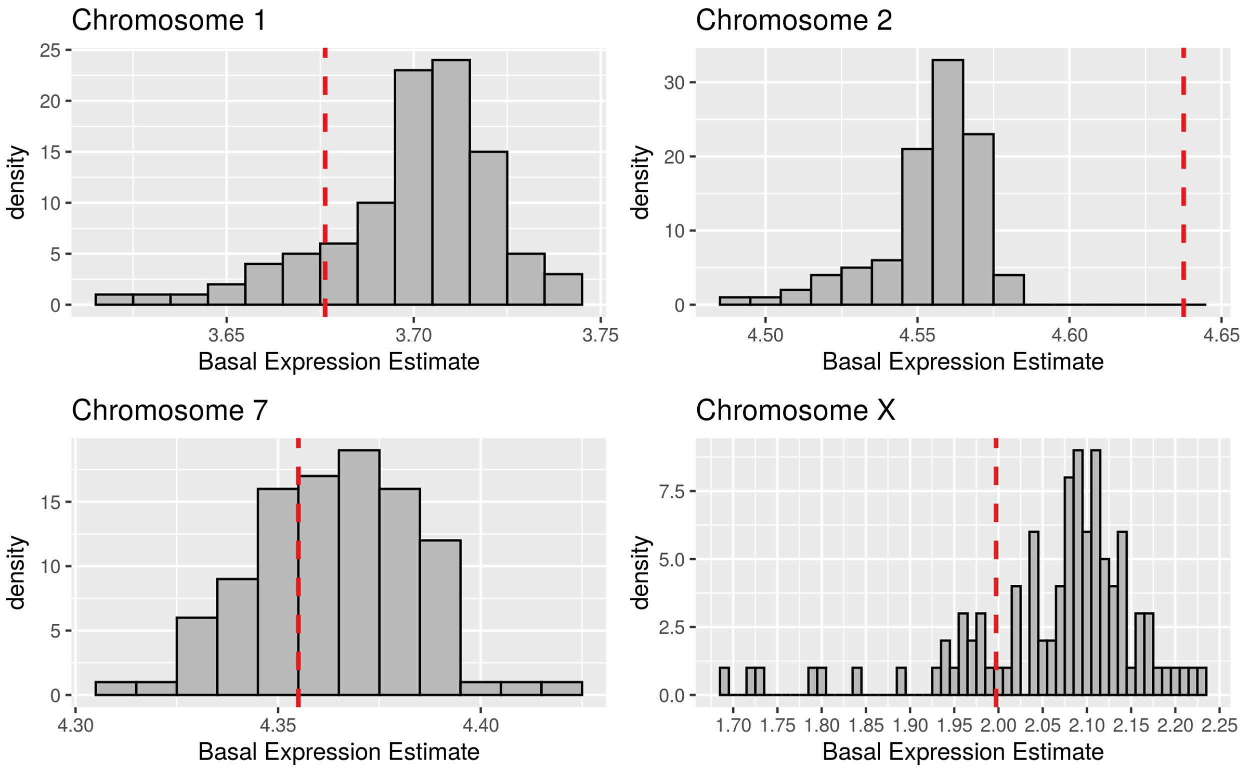

Figure S5: **Histogram of network properties for 23 intra-chromosomal gene networks versus 253 inter-chromosomal gene networks. Left:** Network density. **Right:** Edge to Node ratio

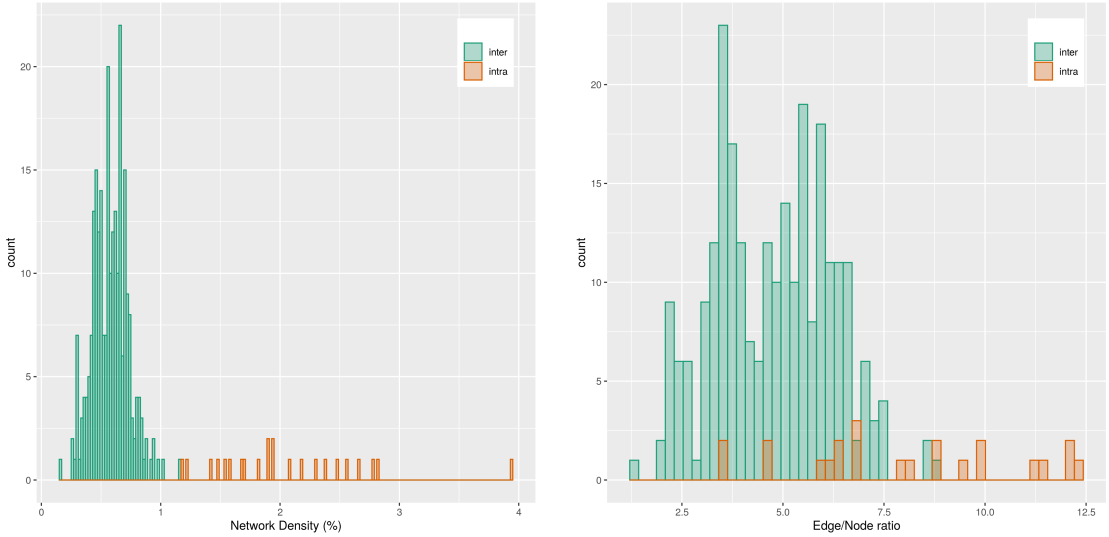

Figure S6: **Differential Expression analysis on GM12878 and IMR90 cell lines.** DE analysis was done using the DESeq2[1] package.Red: genes that are differentially expressed. Green: genes that are not differentially expressed.

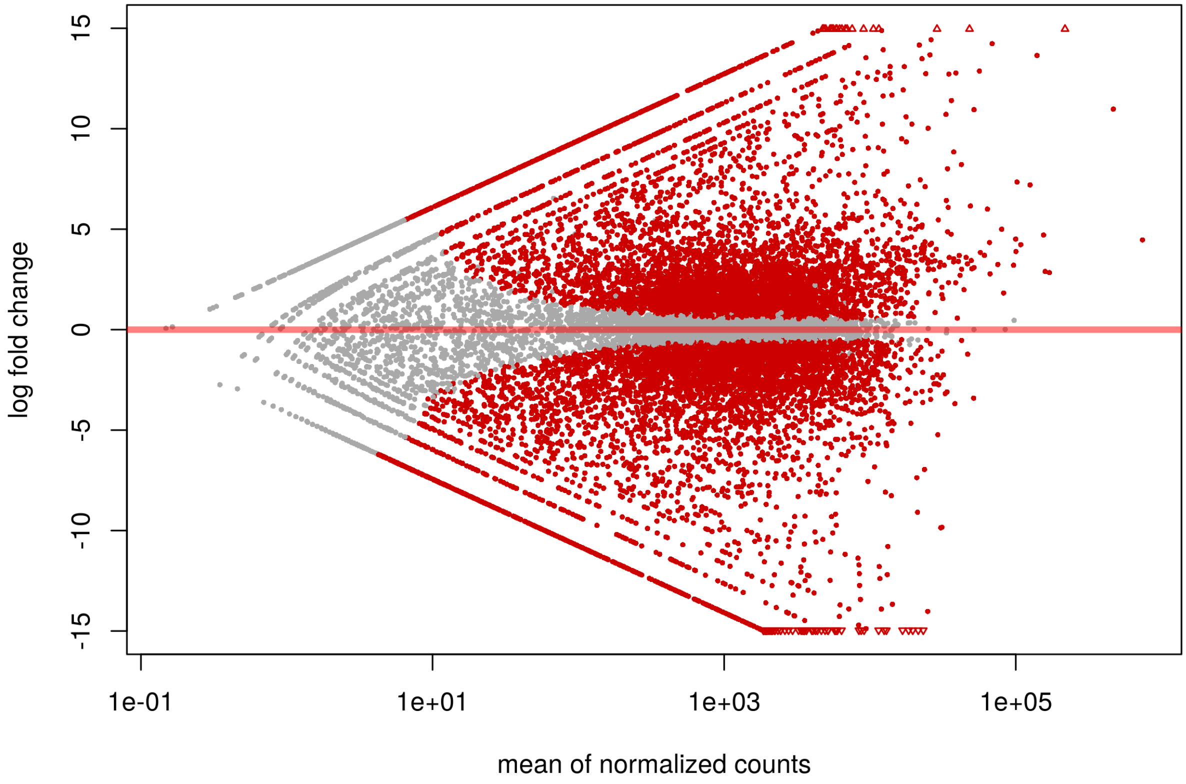

Figure S7: **GM12878: Spatial Interaction Estimate (SIE) for whole chromosomes.** Top panels depicts the network properties of each chromosome. Network density is defined as the percentage of actual edges versus number of possible edges. **(a)** Chromosomes with 95% credible interval above zero. Height of bar is SIE, and error bars are the 2.5% and 97.5% quantiles from the posterior distributions. **(b)** Chromosomes with 95% credible interval including 0. Error bars are the 2.5% and 97.5% quantiles from the posterior distributions.

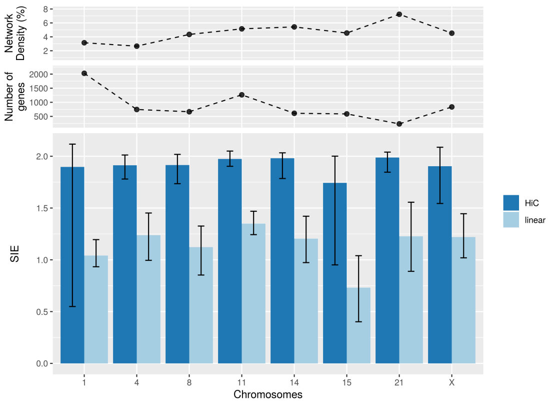

(a)

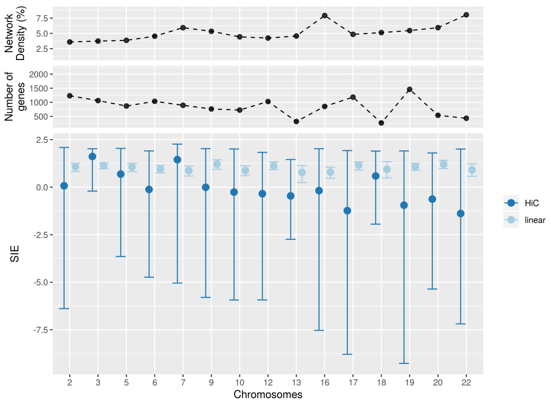

(b)

Figure S8: **GM12878: SIE of chromosome pairs using only inter-chromosomal HiC interactions.** For each pair of chromosomes, PhiMRF ran on a dataset where all genes in both chromosomes are included, while only inter-chromosomal edges are included. From left to right, chromosome pairs ranked by highest SIE to lowest. **Background:** SIE and 95% credible interval of all 253 chromosome pairs. **Zoomed overlay:** SIE and 95% credible interval of the top ten chromosome pairs with the highest SIE.

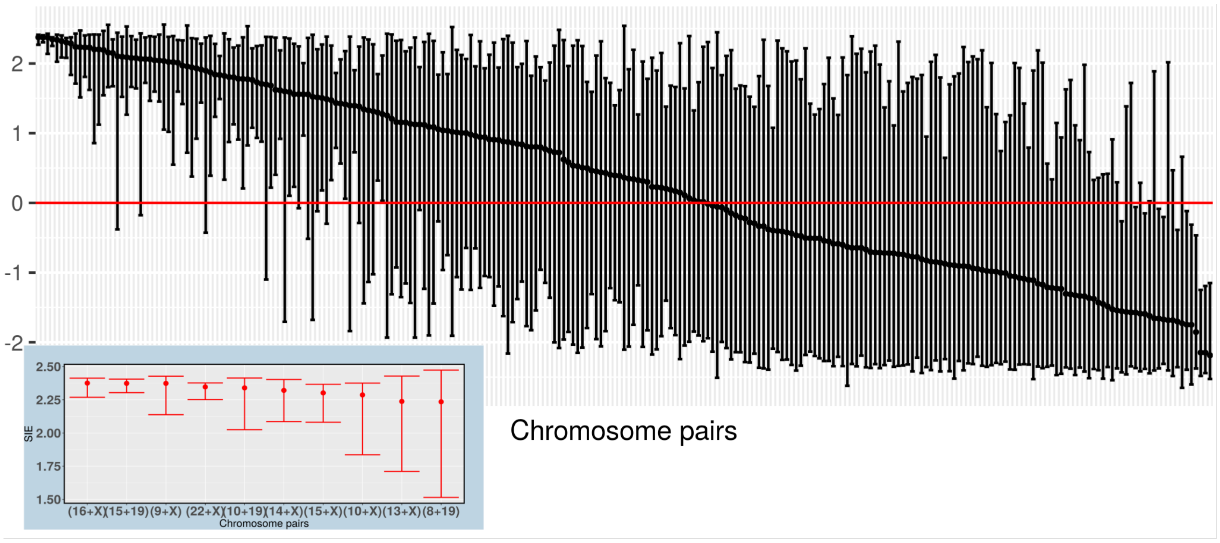

Table S1: **Posterior parameter estimates for all 23 chromosomes** All genes and all edges within each chromosome used.

| Chromosomes | $\hat{\eta}$ | $\hat{\alpha}$ | $\hat{\tau 2}$ |
|---|---|---|---|
| Chrm 1 | 2.060 ( 1.972, 2.108) | 3.490 ( 2.879, 4.185) | 19.176 (16.758, 21.849) |
| Chrm 2 | 1.612 (-0.032, 2.035) | 4.304 ( 3.636, 4.922) | 18.080 (15.474, 20.871) |
| Chrm 3 | 0.930 (-0.639, 1.648) | 4.306 ( 3.665, 4.777) | 20.553 (17.685, 23.287) |
| Chrm 4 | 1.703 ( 1.360, 1.902) | 2.261 ( 1.012, 3.479) | 21.432 (17.121, 27.478) |
| Chrm 5 | 1.298 ( 0.195, 1.874) | 4.458 ( 3.773, 5.023) | 17.601 (14.376, 20.813) |
| Chrm 6 | 1.541 ( 0.530, 1.965) | 3.927 ( 2.935, 4.818) | 19.483 (15.718, 24.077) |
| Chrm 7 | 0.866 (-4.277, 2.184) | 4.194 ( 3.549, 4.793) | 18.893 (13.457, 22.366) |
| Chrm 8 | 1.714 ( 1.395, 1.954) | 2.107 ( 0.458, 3.389) | 21.119 (16.314, 27.164) |
| Chrm 9 | 1.709 ( 1.455, 1.927) | 2.983 ( 2.037, 3.899) | 18.520 (14.447, 24.265) |
| Chrm 10 | 1.019 (-0.867, 1.829) | 4.137 ( 3.208, 4.741) | 19.360 (16.401, 22.535) |
| Chrm 11 | 1.673 (-0.281, 2.049) | 3.539 ( 2.450, 4.551) | 21.877 (18.270, 26.209) |
| Chrm 12 | 1.484 ( 0.561, 1.897) | 4.024 ( 3.119, 4.767) | 20.372 (16.953, 24.526) |
| Chrm 13 | -0.055 (-1.650, 1.194) | 4.443 ( 3.757, 5.038) | 20.128 (15.983, 24.741) |
| Chrm 14 | 1.157 (-1.829, 1.941) | 4.054 ( 2.617, 4.993) | 18.014 (14.151, 22.605) |
| Chrm 15 | 0.770 (-1.497, 1.910) | 4.422 ( 3.185, 5.020) | 17.823 (14.651, 21.681) |
| Chrm 16 | 0.623 (-3.943, 1.980) | 4.011 ( 3.096, 4.580) | 16.745 (13.117, 20.321) |
| Chrm 17 | 1.640 (-0.904, 2.030) | 3.898 ( 3.252, 4.548) | 18.427 (15.367, 21.688) |
| Chrm 18 | 0.335 (-1.177, 1.381) | 4.224 ( 3.293, 4.963) | 23.511 (18.839, 29.021) |
| Chrm 19 | 1.789 ( 1.040, 1.965) | 2.500 ( 0.860, 3.884) | 15.874 (13.240, 19.101) |
| Chrm 20 | 1.376 ( 0.653, 1.873) | 3.177 ( 1.952, 4.065) | 25.312 (20.234, 30.899) |
| Chrm 21 | 1.921 ( 1.712, 1.999) | 0.551 (-1.118, 1.986) | 20.319 (12.405, 30.786) |
| Chrm 22 | 0.565 (-2.582, 1.849) | 4.155 ( 2.878, 4.864) | 17.140 (13.585, 21.259) |
| Chrm X | 1.521 ( 0.894, 2.008) | 1.791 ( 0.806, 2.690) | 31.338 (26.559, 36.224) |

Table S2: **Posterior parameter estimates for the linear baseline** All genes and only linear edges within each chromosome used.

| Chromosomes | $\hat{\eta}$ | $\hat{\alpha}$ | $\hat{\tau^2}$ |
|---|---|---|---|
| Chrm 1 | 1.096 (0.977, 1.256) | 3.297 (2.971, 3.627) | 19.946 (17.864, 22.272) |
| Chrm 2 | 0.850 (0.620, 1.070) | 4.405 (4.095, 4.738) | 18.111 (16.182, 20.164) |
| Chrm 3 | 0.940 (0.722, 1.127) | 4.039 (3.647, 4.428) | 18.010 (15.582, 20.531) |
| Chrm 4 | 1.149 (0.874, 1.424) | 3.390 (2.887, 3.825) | 20.370 (17.145, 24.308) |
| Chrm 5 | 0.958 (0.662, 1.236) | 4.550 (4.172, 4.924) | 16.153 (14.000, 19.035) |
| Chrm 6 | 1.152 (0.922, 1.339) | 3.715 (3.358, 4.078) | 18.399 (15.947, 21.487) |
| Chrm 7 | 0.680 (0.390, 0.963) | 3.930 (3.579, 4.318) | 19.425 (17.187, 22.290) |
| Chrm 8 | 1.227 (0.990, 1.459) | 3.940 (3.481, 4.365) | 17.083 (14.163, 20.957) |
| Chrm 9 | 1.049 (0.758, 1.270) | 3.932 (3.522, 4.316) | 17.684 (15.055, 20.839) |
| Chrm 10 | 0.851 (0.554, 1.150) | 4.189 (3.760, 4.590) | 17.654 (15.305, 20.590) |
| Chrm 11 | 1.276 (1.128, 1.410) | 2.137 (1.649, 2.634) | 22.052 (18.655, 25.978) |
| Chrm 12 | 1.225 (1.032, 1.385) | 3.831 (3.429, 4.258) | 17.062 (14.381, 20.630) |
| Chrm 13 | 0.750 (0.207, 1.185) | 4.271 (3.652, 4.842) | 19.060 (15.359, 23.777) |
| Chrm 14 | 1.058 (0.755, 1.316) | 4.058 (3.543, 4.555) | 16.313 (13.502, 19.772) |
| Chrm 15 | 0.672 (0.321, 1.022) | 4.475 (4.061, 4.888) | 16.874 (14.366, 19.848) |
| Chrm 16 | 0.677 (0.370, 1.004) | 3.886 (3.482, 4.272) | 16.261 (14.230, 18.503) |
| Chrm 17 | 1.081 (0.865, 1.257) | 3.409 (2.995, 3.824) | 17.672 (15.398, 20.266) |
| Chrm 18 | 0.929 (0.425, 1.369) | 3.822 (3.072, 4.489) | 20.829 (15.828, 27.252) |
| Chrm 19 | 1.011 (0.792, 1.230) | 3.218 (2.859, 3.525) | 15.764 (13.838, 18.060) |
| Chrm 20 | 1.150 (0.856, 1.438) | 2.856 (2.203, 3.478) | 21.466 (17.471, 26.499) |
| Chrm 21 | 1.383 (0.998, 1.712) | 2.001 (0.724, 3.106) | 30.121 (19.873, 46.616) |
| Chrm 22 | 0.841 (0.426, 1.200) | 4.010 (3.415, 4.516) | 14.718 (12.042, 18.159) |
| Chrm X | 1.093 (0.809, 1.304) | 1.993 (1.492, 2.480) | 27.328 (23.241, 31.928) |

Table S3: **Summary statistics of the counts of genes and TADs for each chromosome.**

| Chrm | Total Genes | % genes in TADs | Total TADs | % TADs with genes | Median genes per TAD | Max genes per TAD | % DE genes |
|---|---|---|---|---|---|---|---|
| 1 | 2028 | 68.24 | 688 | 50.15 | 3 | 36 | 54.24 |
| 2 | 1232 | 65.91 | 628 | 43.79 | 2 | 19 | 59.98 |
| 3 | 1061 | 66.82 | 549 | 40.98 | 2 | 22 | 56.74 |
| 4 | 745 | 61.74 | 456 | 37.5 | 2 | 12 | 50.47 |
| 5 | 868 | 65.78 | 457 | 37.86 | 2 | 20 | 56.34 |
| 6 | 1035 | 64.25 | 461 | 41 | 2 | 28 | 57.78 |
| 7 | 894 | 57.61 | 411 | 43.31 | 2 | 14 | 55.59 |
| 8 | 665 | 62.56 | 381 | 36.75 | 2 | 19 | 55.64 |
| 9 | 763 | 59.5 | 321 | 37.07 | 2 | 33 | 56.36 |
| 10 | 725 | 66.62 | 366 | 37.7 | 2 | 27 | 57.79 |
| 11 | 1265 | 72.81 | 392 | 49.74 | 3 | 35 | 49.8 |
| 12 | 1028 | 69.46 | 398 | 43.97 | 3 | 22 | 58.17 |
| 13 | 319 | 72.41 | 223 | 35.43 | 2 | 7 | 57.05 |
| 14 | 610 | 68.85 | 249 | 42.97 | 3 | 35 | 55.9 |
| 15 | 588 | 65.65 | 277 | 43.32 | 2 | 16 | 58.67 |
| 16 | 853 | 60.73 | 194 | 56.7 | 3 | 29 | 56.62 |
| 17 | 1183 | 66.95 | 278 | 52.88 | 4 | 30 | 53.51 |
| 18 | 268 | 51.87 | 185 | 30.81 | 1 | 20 | 54.48 |
| 19 | 1459 | 73.06 | 179 | 64.25 | 7 | 43 | 54.83 |
| 20 | 539 | 62.89 | 189 | 43.92 | 2 | 35 | 53.43 |
| 21 | 234 | 65.81 | 86 | 39.53 | 2.5 | 20 | 44.44 |
| 22 | 434 | 65.9 | 118 | 49.15 | 4 | 19 | 58.06 |
| X | 835 | 40.48 | 194 | 53.09 | 2 | 29 | 46.71 |

Table S4: **Posterior parameter estimates for pairwise inter-chromosomal networks** For each pair of chromosomes, all genes from both chromosomes and only inter-chromosomal edges are used.

| Chromosomes | $\hat{\eta}$ | $\hat{\alpha}$ | $\hat{\tau^2}$ |
|---|---|---|---|
| Chrm 1+Chrm 2 | 2.057 ( 1.875, 2.280) | 2.974 (2.707, 3.277) | 22.340 (20.152, 24.862) |
| Chrm 1+Chrm 3 | 2.130 ( 1.802, 2.339) | 3.026 (2.747, 3.353) | 22.652 (20.820, 25.275) |
| Chrm 1+Chrm 4 | 2.194 ( 1.859, 2.400) | 3.035 (2.737, 3.277) | 23.293 (21.278, 25.372) |
| Chrm 1+Chrm 5 | 2.019 ( 1.684, 2.314) | 3.267 (2.941, 3.513) | 22.062 (20.120, 24.298) |
| Chrm 1+Chrm 6 | 2.194 ( 1.803, 2.412) | 3.293 (3.015, 3.554) | 22.724 (21.112, 24.788) |
| Chrm 1+Chrm 7 | 1.956 ( 1.826, 1.999) | 3.165 (2.914, 3.468) | 21.063 (19.323, 22.780) |
| Chrm 1+Chrm 8 | 2.181 ( 1.828, 2.449) | 3.067 (2.761, 3.378) | 22.811 (20.748, 25.290) |
| Chrm 1+Chrm 9 | 1.937 ( 1.709, 2.157) | 3.105 (2.847, 3.376) | 22.697 (20.838, 24.678) |
| Chrm 1+Chrm 10 | 2.053 ( 1.754, 2.289) | 3.022 (2.748, 3.287) | 22.351 (20.081, 24.897) |
| Chrm 1+Chrm 11 | 2.271 ( 2.057, 2.403) | 2.537 (2.220, 2.839) | 23.983 (22.102, 26.363) |
| Chrm 1+Chrm 12 | 2.086 ( 1.776, 2.345) | 3.237 (2.934, 3.561) | 22.712 (20.844, 24.839) |
| Chrm 1+Chrm 13 | 1.842 ( 1.196, 2.317) | 3.330 (3.043, 3.644) | 23.415 (21.328, 25.502) |
| Chrm 1+Chrm 14 | 2.125 ( 1.763, 2.386) | 3.072 (2.804, 3.349) | 22.396 (20.235, 24.861) |
| Chrm 1+Chrm 15 | 2.324 ( 1.969, 2.499) | 3.072 (2.807, 3.306) | 21.675 (19.371, 24.417) |
| Chrm 1+Chrm 16 | 2.291 ( 2.078, 2.433) | 2.974 (2.721, 3.188) | 20.971 (19.079, 22.792) |
| Chrm 1+Chrm 17 | 2.283 ( 2.044, 2.423) | 2.937 (2.691, 3.159) | 21.760 (20.106, 23.794) |
| Chrm 1+Chrm 18 | 2.014 ( 1.451, 2.555) | 3.422 (3.136, 3.743) | 23.741 (21.751, 26.124) |
| Chrm 1+Chrm 19 | 2.224 ( 1.985, 2.340) | 3.007 (2.691, 3.262) | 20.511 (19.089, 22.355) |
| Chrm 1+Chrm 20 | 2.200 ( 1.721, 2.494) | 3.156 (2.860, 3.427) | 23.685 (21.693, 25.869) |
| Chrm 1+Chrm 21 | 1.160 ( 0.081, 1.945) | 3.399 (3.130, 3.636) | 25.369 (23.240, 27.371) |
| Chrm 1+Chrm 22 | 2.147 ( 1.670, 2.537) | 3.285 (3.000, 3.604) | 22.080 (20.444, 23.998) |
| Chrm 1+Chrm X | 1.506 ( 0.667, 1.989) | 3.003 (2.717, 3.296) | 26.000 (24.082, 27.978) |
| Chrm 2+Chrm 3 | 1.892 ( 1.622, 1.999) | 3.885 (3.581, 4.190) | 19.208 (17.424, 21.165) |
| Chrm 2+Chrm 4 | 2.214 ( 1.855, 2.351) | 3.413 (3.010, 3.751) | 19.846 (17.713, 22.222) |
| Chrm 2+Chrm 5 | 1.990 ( 1.487, 2.306) | 4.017 (3.650, 4.416) | 18.703 (16.838, 20.785) |
| Chrm 2+Chrm 6 | 2.145 ( 1.382, 2.355) | 4.002 (3.647, 4.330) | 19.209 (17.420, 21.292) |
| Chrm 2+Chrm 7 | 2.174 ( 1.702, 2.354) | 3.695 (3.347, 4.049) | 19.419 (17.608, 21.473) |
| Chrm 2+Chrm 8 | 2.122 ( 1.684, 2.368) | 3.800 (3.467, 4.154) | 19.484 (17.415, 21.927) |
| Chrm 2+Chrm 9 | 1.958 ( 1.708, 2.193) | 3.537 (3.276, 3.839) | 19.870 (17.807, 22.554) |
| Chrm 2+Chrm 10 | 2.134 ( 1.857, 2.335) | 3.629 (3.333, 3.956) | 19.145 (16.922, 21.554) |
| Chrm 2+Chrm 11 | 2.347 ( 2.229, 2.382) | 2.745 (2.403, 3.081) | 20.950 (18.817, 23.220) |
| Chrm 2+Chrm 12 | 1.900 ( 1.498, 1.996) | 3.965 (3.657, 4.301) | 19.194 (17.571, 21.422) |
| Chrm 2+Chrm 13 | 2.155 ( 1.686, 2.473) | 3.977 (3.656, 4.309) | 19.378 (17.029, 21.709) |
| Chrm 2+Chrm 14 | 1.744 ( 1.328, 1.977) | 3.937 (3.615, 4.261) | 18.844 (16.707, 20.660) |
| Chrm 2+Chrm 15 | 2.098 ( 1.609, 2.400) | 3.895 (3.517, 4.235) | 19.126 (16.811, 21.720) |
| Chrm 2+Chrm 16 | 2.190 ( 1.896, 2.351) | 3.629 (3.314, 3.980) | 18.023 (16.099, 20.296) |
| Chrm 2+Chrm 17 | 2.309 ( 2.058, 2.406) | 3.511 (3.185, 3.807) | 19.061 (17.189, 21.096) |
| Chrm 2+Chrm 18 | 1.482 ( 0.277, 2.362) | 4.183 (3.815, 4.503) | 20.685 (18.785, 22.733) |
| Chrm 2+Chrm 19 | 2.279 ( 1.925, 2.361) | 3.402 (3.100, 3.680) | 17.937 (16.406, 19.799) |
| Chrm 2+Chrm 20 | 1.746 ( 0.403, 2.405) | 3.907 (3.516, 4.271) | 21.231 (19.212, 23.464) |
| Chrm 2+Chrm 21 | 1.287 ( 0.051, 1.950) | 4.016 (3.697, 4.273) | 22.227 (20.238, 24.206) |
| Chrm 2+Chrm 22 | 1.575 ( 0.670, 1.988) | 4.136 (3.767, 4.502) | 18.922 (17.055, 20.613) |
| Chrm 2+Chrm X | 1.908 ( 1.490, 1.999) | 3.131 (2.827, 3.495) | 22.635 (20.372, 24.571) |

| Chromosomes | $\hat{\eta}$ | $\hat{\alpha}$ | $\hat{\tau^2}$ |
|---|---|---|---|
| Chrm 3+Chrm 4 | 2.085 ( 1.673, 2.312) | 3.428 (2.983, 3.833) | 21.139 (18.879, 23.662) |
| Chrm 3+Chrm 5 | 1.235 (-0.260, 1.981) | 4.325 (3.970, 4.675) | 19.861 (18.131, 21.885) |
| Chrm 3+Chrm 6 | 1.830 ( 0.360, 2.344) | 4.024 (3.665, 4.345) | 20.796 (18.698, 23.000) |
| Chrm 3+Chrm 7 | 1.782 ( 1.002, 1.992) | 3.808 (3.425, 4.273) | 19.490 (17.839, 21.615) |
| Chrm 3+Chrm 8 | 1.815 ( 1.436, 1.988) | 3.871 (3.525, 4.241) | 19.811 (17.967, 21.974) |
| Chrm 3+Chrm 9 | 1.616 ( 1.136, 2.077) | 3.740 (3.352, 4.122) | 20.663 (18.422, 23.072) |
| Chrm 3+Chrm 10 | 2.022 ( 1.582, 2.325) | 3.663 (3.298, 4.089) | 20.171 (17.755, 22.872) |
| Chrm 3+Chrm 11 | 2.310 ( 2.147, 2.378) | 2.681 (2.369, 3.034) | 22.395 (20.314, 24.649) |
| Chrm 3+Chrm 12 | 0.714 (-1.159, 2.042) | 4.252 (4.003, 4.501) | 21.572 (19.894, 23.189) |
| Chrm 3+Chrm 13 | 1.583 ( 0.914, 2.198) | 4.030 (3.718, 4.377) | 20.539 (18.505, 23.185) |
| Chrm 3+Chrm 14 | 1.850 ( 1.455, 2.254) | 3.830 (3.415, 4.155) | 19.996 (17.928, 22.882) |
| Chrm 3+Chrm 15 | 1.866 ( 1.380, 2.284) | 3.996 (3.564, 4.347) | 19.571 (17.412, 22.361) |
| Chrm 3+Chrm 16 | 2.152 ( 1.823, 2.329) | 3.590 (3.237, 3.943) | 18.381 (16.299, 20.897) |
| Chrm 3+Chrm 17 | 2.198 ( 1.802, 2.384) | 3.511 (3.212, 3.806) | 19.860 (17.936, 22.305) |
| Chrm 3+Chrm 18 | 0.781 (-0.660, 1.851) | 4.322 (4.016, 4.578) | 21.295 (19.404, 23.740) |
| Chrm 3+Chrm 19 | 2.288 ( 2.079, 2.372) | 3.330 (3.021, 3.633) | 18.036 (16.301, 20.233) |
| Chrm 3+Chrm 20 | 1.848 ( 0.758, 2.346) | 3.825 (3.468, 4.201) | 21.678 (19.424, 24.214) |
| Chrm 3+Chrm 21 | 0.304 (-1.247, 1.931) | 3.992 (3.697, 4.280) | 23.862 (21.584, 26.520) |
| Chrm 3+Chrm 22 | 1.442 ( 0.268, 1.956) | 4.209 (3.839, 4.543) | 19.314 (17.376, 21.153) |
| Chrm 3+Chrm X | 1.858 ( 1.449, 1.996) | 2.982 (2.600, 3.376) | 24.139 (21.895, 26.779) |
| Chrm 4+Chrm 5 | 1.586 ( 1.078, 1.945) | 3.714 (3.329, 4.094) | 20.398 (18.184, 23.179) |
| Chrm 4+Chrm 6 | 1.805 ( 0.644, 2.294) | 3.490 (3.083, 4.023) | 22.679 (20.796, 25.191) |
| Chrm 4+Chrm 7 | 1.896 ( 1.312, 2.256) | 3.196 (2.741, 3.701) | 22.083 (19.938, 24.665) |
| Chrm 4+Chrm 8 | 1.927 ( 1.674, 2.223) | 2.907 (2.467, 3.298) | 22.157 (19.763, 24.882) |
| Chrm 4+Chrm 9 | 1.790 ( 1.553, 2.126) | 2.742 (2.325, 3.117) | 21.562 (19.079, 24.822) |
| Chrm 4+Chrm 10 | 1.982 ( 1.743, 2.227) | 2.736 (2.256, 3.245) | 21.621 (19.041, 25.037) |
| Chrm 4+Chrm 11 | 2.207 ( 1.929, 2.368) | 2.065 (1.657, 2.442) | 25.344 (22.877, 28.471) |
| Chrm 4+Chrm 12 | 1.667 ( 1.000, 2.113) | 3.558 (3.170, 3.883) | 22.159 (20.050, 24.497) |
| Chrm 4+Chrm 13 | 1.844 ( 1.425, 2.197) | 2.890 (2.392, 3.355) | 21.937 (19.049, 25.394) |
| Chrm 4+Chrm 14 | 1.807 ( 1.402, 2.091) | 2.955 (2.422, 3.450) | 21.433 (18.774, 25.234) |
| Chrm 4+Chrm 15 | 1.552 ( 0.962, 2.024) | 3.469 (2.928, 3.883) | 20.980 (18.723, 23.644) |
| Chrm 4+Chrm 16 | 2.028 ( 1.656, 2.328) | 3.031 (2.587, 3.397) | 19.922 (17.736, 22.908) |
| Chrm 4+Chrm 17 | 2.122 ( 1.756, 2.375) | 2.992 (2.656, 3.309) | 21.363 (19.029, 24.219) |
| Chrm 4+Chrm 18 | 1.947 ( 1.515, 2.317) | 3.027 (2.505, 3.498) | 23.031 (19.910, 26.466) |
| Chrm 4+Chrm 19 | 2.116 ( 1.601, 2.407) | 2.983 (2.625, 3.298) | 20.028 (17.866, 22.449) |
| Chrm 4+Chrm 20 | 1.768 ( 1.275, 1.997) | 2.760 (2.356, 3.180) | 23.882 (21.241, 26.501) |
| Chrm 4+Chrm 21 | 1.833 ( 1.199, 2.307) | 2.719 (2.236, 3.091) | 25.836 (22.618, 29.322) |
| Chrm 4+Chrm 22 | 1.979 ( 1.525, 2.292) | 3.169 (2.753, 3.589) | 20.458 (17.812, 23.820) |
| Chrm 4+Chrm X | 2.043 ( 1.756, 2.286) | 1.605 (1.173, 2.055) | 27.288 (24.227, 30.518) |

| Chromosomes | $\hat{\eta}$ | $\hat{\alpha}$ | $\hat{\tau^2}$ |
|---|---|---|---|
| Chrm 5+Chrm 6 | 0.932 (-1.354, 2.276) | 4.306 (4.002, 4.551) | 20.184 (18.241, 21.915) |
| Chrm 5+Chrm 7 | 2.050 ( 1.269, 2.327) | 3.999 (3.639, 4.402) | 18.260 (16.432, 20.184) |
| Chrm 5+Chrm 8 | 1.407 (-0.048, 2.245) | 4.223 (3.886, 4.580) | 19.484 (17.092, 21.747) |
| Chrm 5+Chrm 9 | 1.777 ( 1.434, 2.096) | 3.771 (3.450, 4.096) | 19.115 (17.036, 21.860) |
| Chrm 5+Chrm 10 | 1.748 ( 0.966, 2.226) | 4.133 (3.802, 4.538) | 18.890 (16.656, 21.042) |
| Chrm 5+Chrm 11 | 2.237 ( 1.910, 2.375) | 2.935 (2.547, 3.283) | 22.347 (20.534, 24.896) |
| Chrm 5+Chrm 12 | 0.978 (-1.407, 2.263) | 4.412 (4.082, 4.730) | 19.788 (17.728, 21.660) |
| Chrm 5+Chrm 13 | 1.425 ( 0.487, 1.963) | 4.378 (3.974, 4.710) | 18.616 (16.813, 20.677) |
| Chrm 5+Chrm 14 | 1.479 ( 0.501, 2.090) | 4.233 (3.864, 4.626) | 18.894 (16.853, 21.421) |
| Chrm 5+Chrm 15 | 1.204 (-0.061, 2.122) | 4.533 (4.143, 4.838) | 18.091 (16.261, 20.197) |
| Chrm 5+Chrm 16 | 1.937 ( 1.039, 2.313) | 4.093 (3.733, 4.498) | 17.287 (15.471, 19.110) |
| Chrm 5+Chrm 17 | 1.872 ( 1.505, 1.996) | 3.765 (3.457, 4.111) | 18.617 (16.839, 20.566) |
| Chrm 5+Chrm 18 | -0.259 (-1.985, 1.468) | 4.594 (4.319, 4.862) | 19.500 (17.754, 21.753) |
| Chrm 5+Chrm 19 | 1.885 ( 0.630, 2.119) | 3.692 (3.442, 3.955) | 17.709 (16.209, 19.700) |
| Chrm 5+Chrm 20 | 1.763 ( 0.451, 2.328) | 4.001 (3.565, 4.332) | 20.589 (18.045, 23.071) |
| Chrm 5+Chrm 21 | 0.866 (-0.825, 2.187) | 4.148 (3.752, 4.487) | 21.937 (19.323, 24.302) |
| Chrm 5+Chrm 22 | 1.079 (-0.400, 2.096) | 4.515 (4.157, 4.805) | 17.922 (16.081, 20.179) |
| Chrm 5+Chrm X | 1.741 ( 1.085, 2.000) | 3.095 (2.735, 3.456) | 23.626 (21.547, 26.111) |
| Chrm 6+Chrm 7 | 1.389 ( 0.354, 1.985) | 3.744 (3.315, 4.089) | 21.992 (20.012, 24.153) |
| Chrm 6+Chrm 8 | 0.759 (-0.427, 1.736) | 3.924 (3.549, 4.221) | 22.286 (20.456, 24.687) |
| Chrm 6+Chrm 9 | 1.245 ( 0.791, 1.671) | 3.637 (3.285, 3.954) | 22.043 (19.856, 24.857) |
| Chrm 6+Chrm 10 | 1.117 ( 0.323, 1.648) | 3.842 (3.495, 4.172) | 21.787 (20.073, 24.099) |
| Chrm 6+Chrm 11 | 1.807 ( 1.238, 2.263) | 2.938 (2.598, 3.294) | 25.326 (23.328, 27.723) |
| Chrm 6+Chrm 12 | 0.525 (-0.861, 1.603) | 4.069 (3.783, 4.303) | 22.479 (20.931, 23.890) |
| Chrm 6+Chrm 13 | 0.470 (-0.722, 1.404) | 3.990 (3.625, 4.317) | 22.198 (20.330, 24.658) |
| Chrm 6+Chrm 14 | 1.481 ( 1.012, 1.849) | 3.637 (3.268, 3.992) | 21.307 (19.066, 23.875) |
| Chrm 6+Chrm 15 | 0.760 (-0.448, 1.639) | 4.105 (3.712, 4.432) | 21.033 (19.216, 23.208) |
| Chrm 6+Chrm 16 | 1.553 ( 0.871, 1.987) | 3.647 (3.277, 4.005) | 20.194 (18.304, 22.700) |
| Chrm 6+Chrm 17 | 1.445 ( 0.765, 1.908) | 3.605 (3.304, 3.863) | 21.899 (20.460, 23.800) |
| Chrm 6+Chrm 18 | -0.175 (-1.811, 1.137) | 4.049 (3.728, 4.316) | 23.364 (21.215, 25.338) |
| Chrm 6+Chrm 19 | 1.209 (-0.231, 2.172) | 3.530 (3.242, 3.802) | 20.826 (19.392, 22.634) |
| Chrm 6+Chrm 20 | 0.867 (-0.235, 1.812) | 3.677 (3.347, 3.992) | 24.229 (22.027, 26.373) |
| Chrm 6+Chrm 21 | 1.003 (-0.195, 1.981) | 3.574 (3.219, 3.913) | 25.360 (22.687, 28.057) |
| Chrm 6+Chrm 22 | 0.849 (-0.299, 1.681) | 4.025 (3.660, 4.331) | 21.069 (19.122, 23.199) |
| Chrm 6+Chrm X | 1.091 ( 0.234, 1.846) | 3.001 (2.660, 3.302) | 26.939 (24.409, 29.326) |

| Chromosomes | $\hat{\eta}$ | $\hat{\alpha}$ | $\hat{\tau^2}$ |
|---|---|---|---|
| Chrm 7+Chrm 8 | 1.365 ( 0.556, 1.962) | 3.658 (3.247, 4.076) | 20.975 (19.154, 23.533) |
| Chrm 7+Chrm 9 | 1.582 ( 1.268, 1.912) | 3.411 (3.062, 3.724) | 20.814 (18.862, 23.499) |
| Chrm 7+Chrm 10 | 1.576 ( 0.992, 1.992) | 3.585 (3.176, 3.979) | 20.351 (18.377, 22.940) |
| Chrm 7+Chrm 11 | 2.124 ( 1.911, 2.359) | 2.407 (2.041, 2.782) | 23.835 (21.426, 26.446) |
| Chrm 7+Chrm 12 | 0.997 (-0.520, 1.943) | 3.985 (3.626, 4.264) | 21.326 (19.783, 23.327) |
| Chrm 7+Chrm 13 | 1.692 ( 1.189, 2.236) | 3.541 (3.197, 3.844) | 20.442 (17.990, 23.671) |
| Chrm 7+Chrm 14 | 1.614 ( 1.150, 2.095) | 3.492 (2.949, 3.860) | 19.980 (17.799, 22.562) |
| Chrm 7+Chrm 15 | 1.588 ( 1.004, 2.085) | 3.700 (3.293, 4.095) | 19.976 (17.952, 22.296) |
| Chrm 7+Chrm 16 | 1.776 ( 1.149, 2.203) | 3.538 (3.198, 3.942) | 18.959 (17.190, 21.321) |
| Chrm 7+Chrm 17 | 1.980 ( 1.493, 2.367) | 3.292 (2.943, 3.612) | 20.619 (18.682, 22.643) |
| Chrm 7+Chrm 18 | 1.360 ( 0.308, 2.107) | 3.768 (3.406, 4.198) | 21.556 (19.257, 24.199) |
| Chrm 7+Chrm 19 | 2.074 ( 1.481, 2.381) | 3.149 (2.871, 3.508) | 19.360 (17.706, 21.370) |
| Chrm 7+Chrm 20 | 1.429 ( 0.328, 2.083) | 3.469 (3.035, 3.858) | 22.924 (20.855, 25.338) |
| Chrm 7+Chrm 21 | 1.216 ( 0.196, 2.084) | 3.413 (3.010, 3.797) | 23.922 (21.242, 26.752) |
| Chrm 7+Chrm 22 | 1.088 (-0.029, 1.875) | 3.936 (3.563, 4.250) | 19.765 (17.838, 22.241) |
| Chrm 7+Chrm X | 2.102 ( 1.788, 2.338) | 2.290 (1.923, 2.672) | 24.889 (22.545, 27.452) |
| Chrm 8+Chrm 9 | 1.717 ( 1.474, 1.964) | 3.293 (2.852, 3.663) | 20.525 (17.985, 23.341) |
| Chrm 8+Chrm 10 | 1.669 ( 1.212, 2.077) | 3.603 (3.139, 3.992) | 20.353 (18.329, 23.058) |
| Chrm 8+Chrm 11 | 2.095 ( 1.726, 2.362) | 2.486 (2.051, 2.923) | 24.909 (22.301, 27.867) |
| Chrm 8+Chrm 12 | 1.418 (-0.046, 2.245) | 3.942 (3.535, 4.315) | 21.535 (18.980, 23.900) |
| Chrm 8+Chrm 13 | 1.685 ( 1.122, 2.137) | 3.534 (2.982, 4.037) | 20.449 (17.543, 24.108) |
| Chrm 8+Chrm 14 | 1.782 ( 1.389, 2.108) | 3.366 (2.939, 3.829) | 20.113 (17.406, 23.488) |
| Chrm 8+Chrm 15 | 1.686 ( 1.088, 2.156) | 3.813 (3.392, 4.287) | 19.232 (17.117, 22.005) |
| Chrm 8+Chrm 16 | 1.732 ( 1.209, 1.983) | 3.618 (3.230, 4.010) | 18.296 (16.377, 20.652) |
| Chrm 8+Chrm 17 | 2.063 ( 1.601, 2.374) | 3.245 (2.858, 3.619) | 20.734 (18.795, 22.955) |
| Chrm 8+Chrm 18 | 1.344 ( 0.276, 2.070) | 3.757 (3.169, 4.256) | 21.986 (19.258, 25.353) |
| Chrm 8+Chrm 19 | 2.199 ( 1.730, 2.430) | 3.121 (2.814, 3.426) | 18.679 (17.023, 20.383) |
| Chrm 8+Chrm 20 | 1.720 ( 1.222, 1.982) | 3.158 (2.670, 3.617) | 22.729 (20.088, 25.285) |
| Chrm 8+Chrm 21 | 1.533 ( 0.774, 1.973) | 3.186 (2.730, 3.657) | 24.376 (21.366, 27.405) |
| Chrm 8+Chrm 22 | 1.443 ( 0.613, 1.912) | 3.872 (3.398, 4.328) | 19.399 (17.406, 21.860) |
| Chrm 8+Chrm X | 1.549 ( 0.902, 2.193) | 2.398 (1.858, 2.884) | 27.070 (24.049, 29.850) |
| Chrm 9+Chrm 10 | 2.180 ( 1.987, 2.275) | 3.114 (2.756, 3.470) | 18.338 (15.749, 21.155) |
| Chrm 9+Chrm 11 | 2.236 ( 2.170, 2.265) | 2.443 (2.131, 2.712) | 21.068 (19.177, 23.206) |
| Chrm 9+Chrm 12 | 2.147 ( 1.874, 2.243) | 3.759 (3.403, 4.086) | 19.077 (17.202, 21.188) |
| Chrm 9+Chrm 13 | 2.358 ( 2.154, 2.465) | 3.155 (2.839, 3.546) | 18.076 (15.066, 21.490) |
| Chrm 9+Chrm 14 | 2.181 ( 1.940, 2.290) | 3.201 (2.817, 3.561) | 18.596 (16.230, 21.496) |
| Chrm 9+Chrm 15 | 2.211 ( 1.918, 2.322) | 3.505 (3.102, 3.880) | 17.559 (15.291, 20.295) |
| Chrm 9+Chrm 16 | 2.180 ( 1.833, 2.281) | 3.585 (3.281, 3.940) | 17.475 (15.596, 19.787) |
| Chrm 9+Chrm 17 | 2.190 ( 1.983, 2.280) | 3.256 (2.972, 3.541) | 18.993 (17.134, 21.335) |
| Chrm 9+Chrm 18 | 1.964 ( 1.817, 1.999) | 3.356 (2.980, 3.707) | 18.027 (15.687, 20.583) |
| Chrm 9+Chrm 19 | 2.210 ( 2.123, 2.238) | 3.163 (2.898, 3.441) | 16.997 (15.596, 18.475) |
| Chrm 9+Chrm 20 | 1.962 ( 1.858, 1.996) | 3.163 (2.799, 3.506) | 19.927 (17.599, 22.460) |
| Chrm 9+Chrm 21 | 2.521 ( 2.135, 2.641) | 3.009 (2.608, 3.424) | 21.328 (18.709, 24.871) |
| Chrm 9+Chrm 22 | 1.929 ( 1.684, 1.998) | 3.692 (3.341, 4.039) | 16.884 (14.812, 19.511) |
| Chrm 9+Chrm X | 2.244 ( 2.155, 2.282) | 2.238 (1.893, 2.599) | 23.090 (20.370, 26.448) |

| Chromosomes | $\hat{\eta}$ | $\hat{\alpha}$ | $\hat{\tau^2}$ |
|---|---|---|---|
| Chrm 10+Chrm 11 | 2.258 ( 2.062, 2.397) | 2.363 (2.015, 2.774) | 23.671 (21.134, 26.703) |
| Chrm 10+Chrm 12 | 2.016 ( 1.282, 2.317) | 3.868 (3.520, 4.270) | 19.797 (17.524, 22.191) |
| Chrm 10+Chrm 13 | 1.850 ( 1.310, 2.306) | 3.704 (3.234, 4.179) | 19.507 (16.658, 22.530) |
| Chrm 10+Chrm 14 | 1.995 ( 1.653, 2.294) | 3.455 (3.005, 3.886) | 18.530 (15.662, 22.135) |
| Chrm 10+Chrm 15 | 1.984 ( 1.638, 2.251) | 3.659 (3.166, 4.170) | 18.634 (16.189, 21.352) |
| Chrm 10+Chrm 16 | 2.133 ( 1.770, 2.339) | 3.367 (2.994, 3.736) | 17.891 (15.672, 20.502) |
| Chrm 10+Chrm 17 | 2.273 ( 2.099, 2.380) | 3.133 (2.806, 3.446) | 19.274 (17.298, 22.102) |
| Chrm 10+Chrm 18 | 1.814 ( 0.930, 2.335) | 3.847 (3.467, 4.274) | 20.310 (17.521, 23.505) |
| Chrm 10+Chrm 19 | 1.971 ( 1.869, 2.000) | 3.203 (2.902, 3.458) | 17.133 (15.612, 18.816) |
| Chrm 10+Chrm 20 | 2.045 ( 1.352, 2.340) | 3.346 (2.905, 3.871) | 21.220 (18.610, 23.969) |
| Chrm 10+Chrm 21 | 1.497 ( 0.490, 1.988) | 3.500 (3.033, 3.963) | 23.188 (20.428, 26.000) |
| Chrm 10+Chrm 22 | 1.904 ( 1.340, 2.281) | 3.777 (3.397, 4.227) | 18.546 (15.625, 21.792) |
| Chrm 10+Chrm X | 2.242 ( 1.957, 2.343) | 1.977 (1.561, 2.401) | 24.028 (21.655, 26.856) |
| Chrm 11+Chrm 12 | 1.795 ( 1.492, 2.215) | 2.803 (2.444, 3.164) | 25.067 (23.091, 27.417) |
| Chrm 11+Chrm 13 | 1.885 ( 1.620, 1.996) | 2.352 (2.004, 2.695) | 25.227 (22.586, 28.318) |
| Chrm 11+Chrm 14 | 1.907 ( 1.643, 2.180) | 2.202 (1.838, 2.579) | 24.770 (22.464, 27.769) |
| Chrm 11+Chrm 15 | 2.034 ( 1.796, 2.230) | 2.269 (1.873, 2.659) | 23.873 (21.337, 27.053) |
| Chrm 11+Chrm 16 | 2.119 ( 1.849, 2.340) | 2.405 (2.018, 2.774) | 23.017 (20.521, 26.297) |
| Chrm 11+Chrm 17 | 1.999 ( 1.713, 2.280) | 2.447 (2.117, 2.797) | 24.112 (22.027, 26.463) |
| Chrm 11+Chrm 18 | 1.331 ( 0.640, 1.933) | 2.727 (2.334, 3.115) | 27.598 (25.204, 30.205) |
| Chrm 11+Chrm 19 | 1.438 ( 0.677, 2.038) | 2.880 (2.488, 3.213) | 22.973 (21.102, 24.909) |
| Chrm 11+Chrm 20 | 1.889 ( 1.462, 2.249) | 2.363 (1.992, 2.741) | 27.453 (25.006, 30.169) |
| Chrm 11+Chrm 21 | 1.749 ( 1.211, 2.310) | 2.371 (2.019, 2.723) | 29.292 (26.415, 32.640) |
| Chrm 11+Chrm 22 | 2.148 ( 1.868, 2.400) | 2.303 (1.915, 2.669) | 24.266 (21.521, 27.893) |
| Chrm 11+Chrm X | 1.837 ( 1.529, 1.998) | 1.871 (1.514, 2.266) | 28.625 (26.293, 31.043) |
| Chrm 12+Chrm 13 | 0.825 (-0.642, 1.784) | 4.096 (3.760, 4.554) | 21.411 (19.299, 23.731) |
| Chrm 12+Chrm 14 | 1.134 ( 0.080, 1.896) | 4.019 (3.672, 4.400) | 21.131 (19.175, 23.415) |
| Chrm 12+Chrm 15 | 0.891 (-0.211, 1.686) | 4.213 (3.855, 4.493) | 20.606 (18.814, 22.558) |
| Chrm 12+Chrm 16 | 1.751 ( 0.851, 2.264) | 3.882 (3.536, 4.228) | 19.567 (17.818, 21.831) |
| Chrm 12+Chrm 17 | 1.723 ( 0.933, 2.234) | 3.695 (3.376, 4.019) | 21.368 (19.370, 23.342) |
| Chrm 12+Chrm 18 | 0.631 (-0.826, 1.730) | 4.126 (3.817, 4.438) | 22.434 (20.492, 24.536) |
| Chrm 12+Chrm 19 | 1.728 (-0.110, 2.310) | 3.567 (3.228, 3.915) | 19.796 (17.939, 21.842) |
| Chrm 12+Chrm 20 | 0.104 (-1.325, 1.405) | 3.912 (3.655, 4.192) | 23.654 (21.470, 25.950) |
| Chrm 12+Chrm 21 | 0.301 (-1.002, 1.597) | 3.796 (3.514, 4.058) | 24.622 (22.186, 27.521) |
| Chrm 12+Chrm 22 | -0.177 (-1.579, 1.100) | 4.254 (4.011, 4.508) | 20.713 (18.812, 22.709) |
| Chrm 12+Chrm X | 1.282 ( 0.279, 1.911) | 3.062 (2.610, 3.426) | 26.013 (23.859, 28.375) |
| Chrm 13+Chrm 14 | 1.673 ( 1.143, 2.185) | 3.786 (3.313, 4.257) | 18.958 (16.213, 22.580) |
| Chrm 13+Chrm 15 | 1.096 (-0.386, 2.033) | 4.336 (3.802, 4.785) | 18.905 (16.481, 21.712) |
| Chrm 13+Chrm 16 | 1.999 ( 1.479, 2.361) | 3.606 (3.209, 4.047) | 17.760 (15.414, 20.212) |
| Chrm 13+Chrm 17 | 1.929 ( 1.497, 2.115) | 3.353 (2.981, 3.707) | 20.115 (18.080, 22.612) |
| Chrm 13+Chrm 18 | 1.745 ( 1.081, 2.193) | 3.442 (2.735, 4.151) | 20.692 (17.254, 25.728) |
| Chrm 13+Chrm 19 | 2.271 ( 1.696, 2.565) | 3.130 (2.878, 3.419) | 18.423 (16.564, 20.227) |
| Chrm 13+Chrm 20 | 1.960 ( 1.510, 2.289) | 2.702 (2.019, 3.338) | 23.029 (19.833, 27.233) |
| Chrm 13+Chrm 21 | 1.783 ( 1.296, 1.984) | 2.598 (1.922, 3.183) | 24.241 (20.338, 28.648) |
| Chrm 13+Chrm 22 | 1.609 ( 0.750, 2.191) | 3.923 (3.420, 4.409) | 17.957 (15.107, 20.954) |
| Chrm 13+Chrm X | 2.099 ( 1.841, 2.346) | 1.498 (0.947, 1.946) | 27.969 (24.711, 31.983) |

| Chromosomes | $\hat{\eta}$ | $\hat{\alpha}$ | $\hat{\tau^2}$ |
|---|---|---|---|
| Chrm 14+Chrm 15 | 2.029 ( 1.552, 2.296) | 3.837 (3.377, 4.245) | 17.839 (15.112, 20.752) |
| Chrm 14+Chrm 16 | 2.125 ( 1.831, 2.309) | 3.532 (3.195, 3.900) | 17.182 (15.022, 19.755) |
| Chrm 14+Chrm 17 | 2.230 ( 1.990, 2.360) | 3.171 (2.810, 3.496) | 19.482 (17.316, 21.763) |
| Chrm 14+Chrm 18 | 1.881 ( 1.100, 2.304) | 3.837 (3.324, 4.331) | 19.489 (16.104, 22.925) |
| Chrm 14+Chrm 19 | 2.279 ( 2.016, 2.408) | 3.167 (2.877, 3.476) | 17.589 (15.917, 19.368) |
| Chrm 14+Chrm 20 | 2.036 ( 1.462, 2.290) | 3.245 (2.794, 3.720) | 21.101 (18.184, 24.359) |
| Chrm 14+Chrm 21 | 1.930 ( 1.068, 2.397) | 3.426 (2.901, 3.930) | 22.558 (19.370, 26.628) |
| Chrm 14+Chrm 22 | 1.832 ( 1.107, 2.298) | 3.961 (3.473, 4.368) | 17.540 (15.422, 20.077) |
| Chrm 14+Chrm X | 2.198 ( 1.985, 2.327) | 1.981 (1.483, 2.398) | 24.540 (21.966, 27.867) |
| Chrm 15+Chrm 16 | 2.152 ( 1.741, 2.330) | 3.716 (3.310, 4.055) | 16.727 (14.445, 18.975) |
| Chrm 15+Chrm 17 | 2.268 ( 1.973, 2.383) | 3.264 (2.949, 3.597) | 19.027 (17.119, 21.376) |
| Chrm 15+Chrm 18 | 1.182 (-0.459, 2.173) | 4.361 (3.925, 4.742) | 19.432 (16.892, 22.501) |
| Chrm 15+Chrm 19 | 2.333 ( 2.132, 2.425) | 3.083 (2.802, 3.361) | 17.380 (15.653, 19.483) |
| Chrm 15+Chrm 20 | 2.023 ( 1.214, 2.327) | 3.448 (3.020, 3.916) | 20.903 (18.010, 24.005) |
| Chrm 15+Chrm 21 | 2.098 ( 1.040, 2.398) | 3.709 (3.215, 4.204) | 20.471 (17.501, 24.099) |
| Chrm 15+Chrm 22 | 1.815 ( 0.793, 2.312) | 4.162 (3.658, 4.584) | 17.053 (14.553, 19.791) |
| Chrm 15+Chrm X | 1.899 ( 1.653, 1.998) | 2.449 (2.038, 2.912) | 23.654 (21.331, 26.908) |
| Chrm 16+Chrm 17 | 1.882 ( 1.491, 2.239) | 3.345 (3.046, 3.677) | 19.317 (17.344, 21.263) |
| Chrm 16+Chrm 18 | 1.692 ( 0.815, 2.230) | 3.719 (3.289, 4.099) | 18.458 (16.212, 21.069) |
| Chrm 16+Chrm 19 | 2.126 ( 1.682, 2.373) | 3.185 (2.886, 3.514) | 18.000 (16.262, 19.826) |
| Chrm 16+Chrm 20 | 1.966 ( 1.253, 2.336) | 3.383 (2.931, 3.759) | 19.852 (18.015, 22.247) |
| Chrm 16+Chrm 21 | 1.907 ( 1.185, 2.420) | 3.329 (2.993, 3.724) | 20.369 (17.847, 22.971) |
| Chrm 16+Chrm 22 | 1.841 ( 1.278, 2.279) | 3.680 (3.298, 4.049) | 17.313 (15.053, 19.913) |
| Chrm 16+Chrm X | 2.031 ( 1.696, 2.292) | 2.324 (1.874, 2.714) | 23.228 (20.581, 25.893) |
| Chrm 17+Chrm 18 | 1.736 ( 1.085, 2.215) | 3.457 (3.166, 3.771) | 21.557 (19.334, 23.996) |
| Chrm 17+Chrm 19 | 1.962 ( 1.364, 2.379) | 3.198 (2.901, 3.503) | 19.832 (18.346, 21.501) |
| Chrm 17+Chrm 20 | 2.003 ( 1.454, 2.333) | 3.166 (2.851, 3.534) | 22.007 (19.962, 24.428) |
| Chrm 17+Chrm 21 | 1.661 ( 0.971, 1.987) | 3.170 (2.885, 3.493) | 22.956 (20.596, 25.054) |
| Chrm 17+Chrm 22 | 1.906 ( 1.474, 2.360) | 3.404 (3.081, 3.698) | 19.898 (17.779, 22.206) |
| Chrm 17+Chrm X | 1.988 ( 1.481, 2.330) | 2.538 (2.222, 2.858) | 24.573 (22.475, 26.929) |
| Chrm 18+Chrm 19 | 2.122 ( 1.353, 2.563) | 3.158 (2.869, 3.465) | 19.508 (17.663, 21.266) |
| Chrm 18+Chrm 20 | 1.683 ( 0.772, 2.255) | 3.102 (2.513, 3.639) | 24.820 (21.882, 28.665) |
| Chrm 18+Chrm 21 | 1.893 ( 1.156, 2.319) | 2.352 (1.588, 3.146) | 28.714 (23.758, 34.876) |
| Chrm 18+Chrm 22 | 1.463 ( 0.484, 1.963) | 3.888 (3.328, 4.438) | 18.629 (16.051, 21.942) |
| Chrm 18+Chrm X | 1.881 ( 1.428, 2.257) | 1.599 (1.075, 2.149) | 30.464 (26.592, 34.515) |
| Chrm 19+Chrm 20 | 1.630 ( 1.032, 2.077) | 2.994 (2.684, 3.304) | 21.221 (19.366, 23.256) |
| Chrm 19+Chrm 21 | 1.103 ( 0.044, 1.905) | 3.046 (2.728, 3.327) | 21.532 (19.492, 23.430) |
| Chrm 19+Chrm 22 | 1.858 ( 1.433, 2.315) | 3.077 (2.792, 3.376) | 18.883 (16.880, 21.176) |
| Chrm 19+Chrm X | 1.095 ( 0.531, 1.637) | 2.717 (2.426, 3.035) | 23.542 (21.635, 25.346) |
| Chrm 20+Chrm 21 | 1.556 ( 0.749, 2.114) | 2.187 (1.540, 2.780) | 30.437 (26.324, 35.457) |
| Chrm 20+Chrm 22 | 1.558 ( 0.805, 2.048) | 3.296 (2.794, 3.809) | 21.985 (19.323, 25.068) |
| Chrm 20+Chrm X | 1.293 ( 0.494, 1.788) | 1.951 (1.462, 2.494) | 30.314 (27.619, 33.297) |
| Chrm 21+Chrm 22 | 1.429 ( 0.707, 1.978) | 3.034 (2.460, 3.631) | 22.821 (19.637, 26.521) |
| Chrm 21+Chrm X | 1.608 ( 1.167, 1.931) | 1.282 (0.726, 1.789) | 32.800 (28.492, 37.857) |
| Chrm 22+Chrm X | 1.631 ( 1.202, 1.960) | 2.208 (1.773, 2.631) | 25.582 (22.876, 28.553) |

Table S5: **Name and counts for top fifteen most annotated GO terms non-propagated (left) versus propagated (right)**. Propagation was carried out for both BPO and MFO terms. Relationships that are propagated through includes: "is-a", "part-of", "regulates", "positively-regulates" and "negatively-regulates".

| No propagation | | | | propagated (including MFO) | | | |
|---|---|---|---|---|---|---|---|
| GO term | GO name | Protein count | Gene count | GO term | GO name | Protein count | Gene count |
| GO:0045944 | positive regulation of transcription by RNA polymerase II | 647 | 504 | GO:0071704 | organic substance metabolic process | 7557 | 5888 |
| GO:0007165 | signal transduction | 572 | 457 | GO:0044237 | cellular metabolic process | 7546 | 5873 |
| GO:0007186 | G protein-coupled receptor signaling pathway | 570 | 491 | GO:0044238 | primary metabolic process | 7163 | 5579 |
| GO:0043312 | neutrophil degranulation | 481 | 373 | GO:0050794 | regulation of cellular process | 7044 | 5498 |
| GO:0000122 | negative regulation of transcription by RNA polymerase II | 412 | 331 | GO:0006807 | nitrogen compound metabolic process | 6810 | 5304 |
| GO:0045893 | positive regulation of transcription, DNA-templated | 389 | 312 | GO:0043170 | macromolecule metabolic process | 6253 | 4901 |
| GO:0043687 | post-translational protein modification | 340 | 264 | GO:0044260 | cellular macromolecule metabolic process | 5507 | 4309 |
| GO:0045892 | negative regulation of transcription, DNA-templated | 332 | 270 | GO:0051716 | cellular response to stimulus | 4910 | 3836 |
| GO:0008284 | positive regulation of cell population proliferation | 315 | 253 | GO:1901564 | organonitrogen compound metabolic process | 4810 | 3739 |
| GO:0043066 | negative regulation of apoptotic process | 297 | 233 | GO:0007154 | cell communication | 4242 | 3336 |
| GO:0006468 | protein phosphorylation | 263 | 203 | GO:0019222 | regulation of metabolic process | 4143 | 3261 |
| GO:0008285 | negative regulation of cell population proliferation | 250 | 208 | GO:0034641 | cellular nitrogen compound metabolic process | 4087 | 3164 |
| GO:0019221 | cytokine-mediated signaling pathway | 239 | 193 | GO:0019538 | protein metabolic process | 4027 | 3149 |
| GO:0010628 | positive regulation of gene expression | 236 | 182 | GO:0007165 | signal transduction | 3972 | 3120 |
| GO:0016579 | protein deubiquitination | 231 | 167 | GO:0009058 | biosynthetic process | 3964 | 3082 |

Table S6: **Posterior parameter estimates for gene groups annotated with top 20 non-propagated BPO terms**

| GO term | $\hat{\eta}$ | $\hat{\alpha}$ | $\hat{\tau^2}$ |
|---|---|---|---|
| GO:0045944 | **1.506 ( 0.732, 1.907)** | 4.254 ( 3.521, 4.928) | 18.647 (15.539, 22.898) |
| GO:0007165 | 1.178 (-0.099, 1.960) | 3.975 ( 3.248, 4.630) | 20.274 (16.801, 24.710) |
| GO:0007186 | **1.360 ( 0.795, 1.875)** | -0.611 (-1.518, 0.288) | 29.510 (23.526, 36.105) |
| GO:0043312 | **1.180 ( 0.023, 1.933)** | 4.972 ( 4.235, 5.687) | 23.662 (18.756, 30.059) |
| GO:0000122 | **1.705 ( 1.248, 1.966)** | 5.406 ( 4.844, 6.001) | 10.700 ( 7.370, 14.256) |
| GO:0045893 | 1.013 (-0.117, 1.705) | 5.199 ( 4.426, 5.861) | 15.353 (12.411, 19.098) |
| GO:0043687 | 0.228 (-1.025, 1.417) | 5.933 ( 5.386, 6.444) | 19.060 (15.450, 23.547) |
| GO:0045892 | 1.221 (-0.144, 2.054) | 5.677 ( 4.927, 6.345) | 13.391 (10.132, 17.539) |
| GO:0008284 | 0.544 (-0.801, 1.610) | 4.963 ( 4.293, 5.621) | 18.553 (14.761, 23.073) |
| GO:0043066 | 0.586 (-0.756, 1.783) | 5.975 ( 5.246, 6.679) | 18.211 (14.625, 22.902) |
| GO:0006468 | 0.711 (-1.015, 1.837) | 6.101 ( 5.568, 6.694) | 10.484 ( 7.896, 13.461) |
| GO:0008285 | **1.235 ( 0.327, 1.875)** | 4.766 ( 3.798, 5.663) | 18.236 (13.948, 23.520) |
| GO:0019221 | 0.304 (-0.435, 0.976) | 2.838 ( 1.816, 3.746) | 33.669 (26.719, 42.229) |
| GO:0010628 | 0.216 (-0.908, 1.186) | 4.338 ( 3.551, 5.086) | 22.805 (18.253, 28.470) |
| GO:0016579 | -0.178 (-1.883, 1.831) | 7.414 ( 6.957, 7.823) | 6.073 ( 3.291, 9.263) |
| GO:0043065 | 0.562 (-1.374, 1.778) | 6.195 ( 5.573, 6.748) | 11.840 ( 8.842, 15.623) |
| GO:0006366 | 0.197 (-1.184, 1.611) | 5.835 ( 5.161, 6.435) | 13.566 (10.796, 16.865) |
| GO:0000165 | 0.554 (-0.497, 1.558) | 4.671 ( 3.861, 5.414) | 22.893 (17.485, 28.794) |
| GO:0000398 | 0.176 (-1.594, 1.715) | 7.899 ( 7.666, 8.188) | 1.876 ( 1.081, 4.232) |
| GO:0016567 | 1.007 (-1.292, 1.875) | 7.433 ( 6.925, 7.931) | 4.478 ( 2.027, 7.557) |

Table S7: **Comparison of intra-chromosomal spatial dependency between IMR90 and GM12878.**

| | | IMR90 | | | GM12878 | | |
|---|---|---|---|---|---|---|---|
| Chrm | Num. DE | Edge count | SIE | Meaningful | Edge count | SIE | Meaningful |
| Chrm 1 | 1100 | 24369 | 2.06 | Yes | 64703 | 1.898 | Yes |
| Chrm 2 | 739 | 10723 | 1.612 | | 27230 | 0.072 | |
| Chrm 3 | 602 | 8373 | 0.93 | | 20983 | 1.614 | |
| Chrm 4 | 376 | 3386 | 1.703 | Yes | 7372 | 1.913 | Yes |
| Chrm 5 | 489 | 5830 | 1.298 | Yes | 14489 | 0.687 | |
| Chrm 6 | 598 | 10169 | 1.541 | Yes | 24249 | -0.12 | |
| Chrm 7 | 497 | 9933 | 0.866 | | 23679 | 1.446 | |
| Chrm 8 | 370 | 4277 | 1.714 | Yes | 9595 | 1.914 | Yes |
| Chrm 9 | 430 | 5283 | 1.709 | Yes | 15513 | -0.007 | |
| Chrm 10 | 419 | 5002 | 1.019 | | 11617 | -0.259 | |
| Chrm 11 | 630 | 15454 | 1.673 | | 41143 | 1.973 | Yes |
| Chrm 12 | 598 | 9018 | 1.484 | Yes | 22321 | -0.344 | |
| Chrm 13 | 182 | 1105 | -0.055 | | 2314 | -0.459 | |
| Chrm 14 | 341 | 4932 | 1.157 | | 10074 | 1.98 | Yes |
| Chrm 15 | 345 | 3574 | 0.77 | | 7843 | 1.743 | Yes |
| Chrm 16 | 483 | 10243 | 0.623 | | 28791 | -0.18 | |
| Chrm 17 | 633 | 11786 | 1.64 | | 33825 | -1.236 | |
| Chrm 18 | 146 | 919 | 0.335 | | 1834 | 0.589 | |
| Chrm 19 | 800 | 16806 | 1.789 | Yes | 58021 | -0.947 | |
| Chrm 20 | 288 | 3450 | 1.376 | Yes | 8606 | -0.628 | |
| Chrm 21 | 104 | 1073 | 1.921 | Yes | 1974 | 1.988 | Yes |
| Chrm 22 | 252 | 2603 | 0.565 | | 7556 | -1.385 | |
| Chrm X | 390 | 7974 | 1.521 | Yes | 15761 | 1.904 | Yes |

Table S8: **Posterior parameter estimates for simulated data.** True parameter values that generated the data are given to compare the efficacy of the model.

| | $\eta$ | $\alpha$ | $\tau^2$ |
|---|---|---|---|
| TRUE | 0.000 | 2.000 | 1.800 |
| Estimate | -0.052 (-0.969, 0.773) | 1.994 (1.894, 2.087) | 1.979 ( 1.782, 2.213) |
| TRUE | 0.500 | 2.000 | 1.800 |
| Estimate | 0.383 (-0.650, 1.274) | 2.004 (1.901, 2.107) | 1.831 ( 1.648, 2.081) |
| TRUE | -0.500 | 2.000 | 1.800 |
| Estimate | -0.130 (-1.129, 0.856) | 2.001 (1.909, 2.085) | 1.725 ( 1.513, 1.914) |
| TRUE | 0.000 | 3.000 | 1.800 |
| Estimate | -0.560 (-1.617, 0.652) | 3.003 (2.922, 3.080) | 1.759 ( 1.458, 1.998) |
| TRUE | 1.000 | 3.000 | 1.800 |
| Estimate | 0.617 (-0.836, 1.704) | 3.095 (2.974, 3.207) | 1.805 ( 1.522, 2.236) |
| TRUE | -1.000 | 3.000 | 1.800 |
| Estimate | -0.591 (-1.654, 0.707) | 3.006 (2.924, 3.084) | 2.031 ( 1.746, 2.292) |
| TRUE | 0.000 | 2.000 | 9.000 |
| Estimate | 0.095 (-0.474, 0.634) | 2.061 (1.853, 2.261) | 9.393 ( 8.366, 10.552) |
| TRUE | 1.500 | 2.000 | 9.000 |
| Estimate | 1.579 ( 1.155, 1.919) | 1.980 (1.531, 2.436) | 9.021 ( 7.706, 10.435) |
| TRUE | -1.500 | 2.000 | 9.000 |
| Estimate | -1.674 (-1.967, -1.356) | 1.923 (1.780, 2.045) | 8.553 ( 7.362, 9.851) |

Table S9: **Posterior parameter estimates using the median metric**. For each chromosome, when summarizing the pool of interactions for each gene pair, **median** is used as the summary statistic.

| Chromosomes | $\hat{\eta}$ | $\hat{\alpha}$ | $\hat{\tau^2}$ |
|---|---|---|---|
| Chrm 1 | 2.029 ( 1.843, 2.102) | 3.696 ( 3.006, 4.401) | 19.616 (16.979, 22.101) |
| Chrm 2 | 1.570 ( 0.062, 2.024) | 4.364 ( 3.712, 4.956) | 18.026 (15.271, 21.112) |
| Chrm 3 | 0.860 (-0.829, 1.548) | 4.351 ( 3.763, 4.842) | 20.545 (17.645, 23.483) |
| Chrm 4 | 1.591 ( 1.290, 1.862) | 2.619 ( 1.409, 3.613) | 21.521 (17.181, 26.410) |
| Chrm 5 | 1.295 ( 0.235, 1.872) | 4.484 ( 3.838, 5.062) | 17.639 (14.597, 21.082) |
| Chrm 6 | 1.575 ( 0.703, 1.975) | 3.899 ( 2.921, 4.825) | 19.538 (16.109, 24.200) |
| Chrm 7 | 0.662 (-4.673, 2.154) | 4.247 ( 3.657, 4.886) | 18.585 (12.801, 22.304) |
| Chrm 8 | 1.576 ( 1.005, 1.929) | 2.684 ( 1.093, 3.792) | 20.871 (16.712, 26.599) |
| Chrm 9 | 1.600 ( 1.283, 1.875) | 3.431 ( 2.556, 4.193) | 18.773 (14.793, 24.547) |
| Chrm 10 | 1.052 (-1.018, 1.986) | 4.158 ( 3.241, 4.773) | 19.441 (16.352, 22.935) |
| Chrm 11 | 1.734 ( 0.233, 2.049) | 3.574 ( 2.527, 4.619) | 21.922 (18.442, 26.360) |
| Chrm 12 | 1.466 ( 0.590, 1.898) | 4.055 ( 3.115, 4.803) | 20.355 (17.146, 25.367) |
| Chrm 13 | 0.089 (-1.459, 1.124) | 4.429 ( 3.788, 4.965) | 20.101 (15.760, 24.667) |
| Chrm 14 | 0.920 (-2.263, 1.906) | 4.229 ( 2.907, 5.053) | 18.332 (14.422, 22.986) |
| Chrm 15 | 0.586 (-1.744, 1.793) | 4.526 ( 3.648, 5.074) | 17.921 (14.963, 21.514) |
| Chrm 16 | 0.529 (-3.669, 1.959) | 4.078 ( 3.212, 4.646) | 16.828 (12.366, 20.394) |
| Chrm 17 | 1.715 ( 0.707, 1.983) | 3.889 ( 3.216, 4.601) | 18.288 (15.843, 21.407) |
| Chrm 18 | 0.440 (-1.057, 1.392) | 4.192 ( 3.191, 4.991) | 23.228 (18.493, 29.125) |
| Chrm 19 | 1.810 ( 1.366, 1.975) | 2.450 ( 0.824, 3.775) | 16.189 (13.502, 19.532) |
| Chrm 20 | 1.415 ( 0.661, 1.867) | 3.161 ( 2.033, 4.116) | 25.087 (20.031, 30.974) |
| Chrm 21 | 1.916 ( 1.686, 1.999) | 0.578 (-1.021, 2.093) | 20.348 (12.211, 31.025) |
| Chrm 22 | 0.385 (-2.494, 1.695) | 4.257 ( 3.218, 4.866) | 17.030 (13.303, 21.071) |
| Chrm X | 1.491 ( 0.721, 1.980) | 1.856 ( 0.975, 2.694) | 31.598 (26.927, 36.267) |

Table S10: **Posterior parameter estimates using the min metric**. For each chromosome, when summarizing the pool of interactions for each gene pair, **min** is used as the summary statistic.

| Chromosomes | $\hat{\eta}$ | $\hat{\alpha}$ | $\hat{\tau^2}$ |
|---|---|---|---|
| Chrm 1 | 1.851 ( 0.836, 2.110) | 4.480 (3.856, 5.062) | 20.868 (18.305, 23.967) |
| Chrm 2 | 1.075 (-0.540, 1.749) | 4.725 (4.342, 5.171) | 19.230 (16.973, 21.848) |
| Chrm 3 | 0.827 ( 0.001, 1.363) | 4.416 (4.016, 4.808) | 20.599 (18.190, 23.310) |
| Chrm 4 | 0.943 ( 0.414, 1.444) | 3.724 (3.262, 4.238) | 22.697 (19.972, 26.233) |
| Chrm 5 | 0.929 ( 0.187, 1.405) | 4.676 (4.279, 5.045) | 17.977 (15.511, 20.598) |
| Chrm 6 | 1.447 ( 1.107, 1.748) | 4.326 (3.750, 4.891) | 20.684 (17.836, 24.258) |
| Chrm 7 | 0.937 (-1.968, 2.127) | 4.246 (3.776, 4.762) | 20.105 (17.160, 23.121) |
| Chrm 8 | 0.803 ( 0.032, 1.391) | 3.875 (3.290, 4.380) | 21.133 (18.106, 24.448) |
| Chrm 9 | 1.215 ( 0.683, 1.595) | 4.284 (3.787, 4.776) | 19.942 (17.111, 23.307) |
| Chrm 10 | 0.574 (-0.582, 1.298) | 4.382 (3.966, 4.771) | 20.198 (17.812, 22.898) |
| Chrm 11 | 1.688 ( 0.375, 1.980) | 4.244 (3.168, 5.208) | 23.904 (20.232, 29.080) |
| Chrm 12 | 1.521 ( 0.950, 1.889) | 4.203 (3.628, 4.801) | 20.048 (17.087, 23.829) |
| Chrm 13 | -0.005 (-0.879, 0.784) | 4.449 (3.884, 5.007) | 20.292 (16.369, 25.101) |
| Chrm 14 | 1.157 ( 0.252, 1.794) | 4.371 (3.656, 5.000) | 18.563 (15.319, 22.607) |
| Chrm 15 | 0.176 (-1.224, 1.333) | 4.657 (4.197, 5.030) | 18.314 (15.919, 21.404) |
| Chrm 16 | 0.314 (-2.668, 1.811) | 4.242 (3.779, 4.746) | 16.968 (14.010, 19.626) |
| Chrm 17 | 1.335 (-0.657, 1.877) | 4.135 (3.571, 4.736) | 19.598 (16.685, 22.671) |
| Chrm 18 | 0.621 (-0.466, 1.390) | 4.333 (3.643, 5.047) | 23.760 (18.914, 29.863) |
| Chrm 19 | 1.430 (-0.422, 1.854) | 3.598 (2.893, 4.398) | 17.307 (14.884, 20.524) |
| Chrm 20 | 1.238 ( 0.701, 1.718) | 3.399 (2.598, 4.156) | 25.872 (21.233, 31.800) |
| Chrm 21 | 1.883 ( 1.655, 1.998) | 1.947 (0.458, 3.133) | 23.633 (14.581, 35.877) |
| Chrm 22 | 0.759 (-0.547, 1.598) | 4.321 (3.579, 4.893) | 16.999 (13.625, 20.862) |
| Chrm X | 1.228 ( 0.429, 1.758) | 2.490 (1.773, 3.205) | 31.727 (26.931, 36.532) |

Table S11: **Posterior parameter estimates using the max metric**. For each chromosome, when summarizing the pool of interactions for each gene pair, **max** is used as the summary statistic.

| Chromosomes | $\hat{\eta}$ | $\hat{\alpha}$ | $\hat{\tau^2}$ |
|---|---|---|---|
| Chrm 1 | 2.133 ( 2.116, 2.135) | 1.664 ( 0.993, 2.269) | 16.951 (14.720, 19.490) |
| Chrm 2 | 2.043 ( 1.964, 2.093) | 3.057 ( 2.289, 3.933) | 16.542 (13.894, 19.490) |
| Chrm 3 | 2.026 ( 1.843, 2.107) | 2.502 ( 1.302, 3.724) | 19.020 (15.451, 23.226) |
| Chrm 4 | 2.049 ( 1.942, 2.089) | 0.501 (-0.504, 1.538) | 16.572 (13.041, 20.881) |
| Chrm 5 | 1.345 (-2.346, 2.073) | 4.348 ( 3.358, 5.166) | 17.500 (13.724, 21.981) |
| Chrm 6 | 1.362 (-5.484, 2.059) | 3.782 ( 2.510, 4.858) | 17.271 (12.217, 21.770) |
| Chrm 7 | 1.215 (-5.390, 2.184) | 3.782 ( 2.848, 4.615) | 18.076 (12.590, 22.116) |
| Chrm 8 | 2.018 ( 1.897, 2.080) | 0.520 (-0.841, 1.716) | 20.136 (15.806, 25.368) |
| Chrm 9 | 1.879 ( 1.694, 1.998) | 1.911 ( 0.758, 3.052) | 18.039 (13.566, 23.511) |
| Chrm 10 | 1.983 ( 1.429, 2.157) | 2.794 ( 1.515, 4.181) | 18.638 (14.324, 24.436) |
| Chrm 11 | 2.086 ( 2.061, 2.089) | -0.459 (-1.720, 0.684) | 18.267 (15.206, 21.514) |
| Chrm 12 | -0.086 (-5.515, 1.965) | 4.429 ( 3.655, 5.242) | 18.563 ( 6.445, 24.058) |
| Chrm 13 | 0.990 (-3.734, 2.028) | 3.553 ( 1.183, 4.986) | 19.186 (14.000, 25.289) |
| Chrm 14 | 2.037 ( 1.927, 2.070) | 0.765 (-0.889, 2.315) | 15.874 (11.832, 20.970) |
| Chrm 15 | 1.278 (-2.976, 2.093) | 3.954 ( 2.524, 5.051) | 17.350 (13.377, 22.693) |
| Chrm 16 | 1.986 ( 1.853, 2.057) | 2.227 ( 1.048, 3.448) | 15.282 (11.988, 19.165) |
| Chrm 17 | 1.528 (-4.079, 2.058) | 3.759 ( 3.012, 4.432) | 18.041 (15.008, 21.780) |
| Chrm 18 | 1.446 (-2.795, 2.077) | 3.126 ( 1.109, 4.911) | 20.587 (14.406, 29.081) |
| Chrm 19 | 1.969 ( 1.917, 2.017) | 0.267 (-1.522, 1.962) | 15.679 (12.908, 19.617) |
| Chrm 20 | 1.826 ( 1.491, 2.025) | 2.363 ( 0.807, 3.616) | 23.761 (19.121, 29.728) |
| Chrm 21 | 1.979 ( 1.835, 2.045) | -0.252 (-1.980, 1.354) | 22.150 (14.298, 31.806) |
| Chrm 22 | 1.045 (-4.053, 1.960) | 3.391 ( 1.296, 4.864) | 16.706 (11.727, 23.081) |
| Chrm X | 2.086 ( 1.946, 2.161) | -1.281 (-2.389, -0.119) | 28.210 (23.897, 32.925) |
